# *Borrelia afzelii* does not suppress the development of anti-tick immunity in bank voles

**DOI:** 10.1101/2020.03.31.018754

**Authors:** Andrea Gomez-Chamorro, Yating Li, Adrian Herrera, Olivier Rais, Hans Dautel, Maarten J. Voordouw

## Abstract

Vector-borne pathogens manipulate their vertebrate hosts to enhance their transmission to arthropod vectors. The ability of vertebrate hosts to develop acquired immunity against arthropod vectors represents an existential threat for both the vector and the pathogen. The purpose of the study was to test whether the tick-borne spirochete bacterium *Borrelia afzelii* could suppress the development of acquired immunity to its tick vector *Ixodes ricinus* in the bank vole *Myodes glareolus*, which is an important host for both the tick and the pathogen. We created a group of *B. afzelii*-infected bank voles and an uninfected control group by exposing lab-reared animals to infected or uninfected ticks. At 1, 2, and 3 months post-infection, all bank voles were infested with larval *I. ricinus* ticks. The bank voles developed a strong antibody response against tick salivary gland extract proteins. This anti-tick immunity had negative effects on tick fitness traits including engorged larval weight, unfed nymphal weight, larva-to-nymph molting time and larva-to-nymph molting success. Infection with *B. afzelii* did not suppress the development of acquired immunity against *I. ricinus* ticks. The development of anti-tick immunity was strongly correlated with a dramatic temporal decline in both the bacterial abundance in the host ear tissues and the host-tick transmission success of *B. afzelii*. Our study suggests that the development of anti-tick immunity in bank voles has important consequences for the density of infected ticks and the risk of Lyme borreliosis.

**Importance:** Many pathogens enhance their persistence and transmission by suppressing the immune system of their host. We used an experimental infection approach to test whether the Lyme disease pathogen, *Borrelia afzelii*, could suppress the development of acquired immunity against its tick vector (*Ixodes ricinus*) in the bank vole (*Myodes glareolus*), but found no evidence for this phenomenon. Uninfected and *B. afzelii*-infected bank voles both developed a strong IgG antibody response against tick salivary gland extract following repeated infestations with *I. ricinus* ticks. The development of anti-tick immunity was negatively correlated with the abundance of *B. afzelii* in ear tissue biopsies and with host-to-tick transmission to *I. ricinus* ticks. Our study suggests that anti-tick immunity in the bank vole reduces the prevalence of this important tick-borne pathogen.

## Introduction

Many pathogens exploit the obligate blood-feeding habits of hematophagous arthropods to achieve transmission between vertebrate hosts. These vector-borne pathogens and their arthropod vectors have co-evolved an intimate relationship that is characterized by specificity and adaptive complexity (1-3). Specificity is shown by the fact that many vector-borne pathogens are only effectively acquired and transmitted by one or a few closely related vector species (4-6). Examples of adaptive complexity in pathogens include increasing the vector’s production of immunosuppressive saliva molecules to enhance their own transmission (7), protecting the vector against changes in the environment (8, 9), and manipulating the odor profile of the vertebrate host to increase host-vector encounter rates and pathogen transmission (10-12). Thus, whenever it suits their interests, we expect vector-borne pathogens to enhance the fitness of their arthropod vectors.

From the perspective of the pathogen and vector, the ability of the vertebrate host to develop immunity or resistance against the vector represents a shared existential threat. Compared to other hematophagous arthropods, this threat of inducing acquired resistance is particularly important for ticks, probably because they feed on the host in large numbers and for longer periods of time (several days). Studies on a variety of host-tick systems have shown that this anti-tick immunity or acquired tick resistance can reduce tick fitness components including measures of larval engorgement, duration of attachment, percentage of recovered larvae, and larva-to-nymph molting success (13-16). Acquired immunity against ticks also enhances the host defense against tick-borne pathogens (17). Acquired immunity against ticks can impair transmission of tick-borne pathogens, establishment of the pathogen within the host, and alter the cutaneous environment at the tick attachment site (17-19). In summary, the development of acquired immunity in the vertebrate host against ticks reduces the fitness of both the tick vector and the tick-borne pathogen.

Ticks and specifically tick salivary glands and tick saliva have evolved to cope with this existential threat of host resistance. Tick saliva contains nearly 500 proteins and peptides that belong to at least 25 different protein families (20). These tick saliva molecules have powerful anti-haemostatic, anti-inflammatory and immunomodulatory properties (21) that seek to subvert the host immune system at every turn. This powerful pharmacopeia also enhances the transmission of pathogens, a phenomenon known as saliva-assisted transmission (22). Conversely, a number of tick-borne pathogens have developed a variety of strategies that allow them to evade or suppress the host immune system (23). In turn, pathogen-induced immunosuppression may reduce the development of anti-tick immunity and thereby enhance tick feeding success.

To study whether vector-borne pathogens can suppress the development of acquired immunity against the arthropod vector in the vertebrate host, we used a tick-borne spirochete bacterium belonging to the *B. burgdorferi* sensu lato (sl) genospecies complex and that causes Lyme borreliosis (LB) in humans as a model system (24, 25). Our study was motivated by a recent demonstration that *B. burgdorferi* sensu stricto (ss) could suppress the development of acquired immunity in the lab mouse *Mus musculus* (26). We wanted to test whether other *B. burgdorferi* sl genospecies could induce immunosuppression in their natural rodent reservoir hosts. The most common LB system in Europe includes the bacterium *B. afzelii*, the tick *Ixodes ricinus*, and rodent hosts such as the bank vole (*Myodes glareolus*) (27-30). The bank vole is a good host for *B. afzelii* (27-29), but it develops acquired immunity against *Ixodes* ticks (14, 27). Previous workers have suggested that this anti-tick immunity in bank voles reduces the production of *B. afzelii*-infected ticks (27).

The purpose of this study was to test whether *B. afzelii* could inhibit the development of acquired anti-tick immunity in bank voles. We created *B. afzelii*-infected bank voles and uninfected bank voles, and repeatedly infested them with larval *I. ricinus* ticks to induce acquired anti-tick immunity. We predicted that bank voles would develop a strong antibody response against tick salivary gland proteins, which would decrease tick fitness. We expected that infection with *B. afzelii* would suppress the development of acquired anti-tick immunity in bank voles. We therefore expected tick fitness to be higher on the *B. afzelii*-infected bank voles compared to the uninfected controls.

## Materials and Methods

### Bank voles, *Ixodes ricinus* ticks and *Borrelia afzelii*

The bank voles came from a laboratory colony at the University of Neuchâtel. This colony was descended from bank voles that had been captured at a field site near Neuchâtel, Switzerland in the summer of 2014 (31). All animals used in the study were lab-born, 5 to 10 weeks old at the start of the study, and not infected with *B. afzelii*. During the experiment, bank voles were maintained in individual cages and were given food and water *ad libitum*. The *I. ricinus* ticks came from the University of Neuchâtel colony, and larval *I. ricinus* ticks were also purchased from Insect Services (Germany). Bank voles were experimentally infected via tick-bite with *B. afzelii* isolate NE4049. This isolate was originally obtained from an *I. ricinus* tick in Neuchâtel, it has multi-locus sequence type 679, *ospC* major group (oMG) A10, and strain ID number 1887 in the *Borrelia* MLST database. We used isolate NE4049 because our previous work has shown that it is highly infectious to *I. ricinus* ticks and rodents including bank voles (31-34). *B. afzelii* strains carrying oMG A10 also have the highest frequency in wild *I. ricinus* populations near Neuchâtel (35).

### Ethics statement and animal experimentation permits

The study was performed at the University of Neuchâtel and followed the Swiss legislation on animal experimentation. The commission that is part of the ‘Service de la Consommation et des Affaires Vétérinaires (SCAV)’ of canton Vaud evaluated and approved the ethics of this part of the study. The SCAV of canton Neuchâtel issued the animal experimentation permits for the study (NE1/2017) and for the maintenance of the *I. ricinus* tick colony at the University of Neuchâtel (NE5/2014).

### Creation of nymphs infected with *B. afzelii*

Larval *I. ricinus* ticks were fed on BALB/c mice (*Mus musculus*) that had been previously infected with *B. afzelii* isolate NE4049 via tick-bite. The resultant engorged larval ticks were collected, stored in individual Eppendorf tubes, and allowed to molt into nymphs. A random sample of 30 nymphs was tested for *B. afzelii* using qPCR, and the prevalence of infection was 93.3% (28 infected/ 30 total). To create uninfected control nymphs, larval *I. ricinus* ticks were fed on uninfected BALB/c mice and the resultant engorged larvae were allowed to molt into nymphs.

### Experimental design

Forty voles were randomly assigned to one of two experimental groups: uninfected control (n = 20) and infected with *B. afzelii* (n = 20). Each vole in the uninfected control group was infested with 5 uninfected nymphs, whereas each vole in the *B. afzelii*-infected group was infested with 5 *B. afzelii*-infected nymphs. During the nymphal challenge, voles were anesthetized with a ketamine/xylazine cocktail via intraperitoneal injection (1:2:9 xylazine: ketamine: PBS; dose of 5 μl per 10 g of vole body mass). Six of the 40 voles died under anesthesia leaving 34 animals. To prevent the voles from removing the nymphs via grooming, the nymphs were placed in a plastic capsule attached to a shaved area on the back of the vole. Each vole was fitted with a collar to prevent the animal from removing the capsule. The capsules were checked on a daily basis and the engorged nymphs were collected and tested for their *B. afzelii* infection status. The infection status of the voles was confirmed using additional diagnostic criteria (see below).

### Larval infestations

Voles were infested with larval ticks on three separate occasions at 27, 54, and 84 days post-infection (PI). For the first and second infestation, we used larvae from our *I. ricinus* tick colony at the University of Neuchâtel. For the third infestation, we had a shortage of larvae at our colony, and we therefore purchased them from Insect Services (Germany). For each infestation, the voles were anaesthetized and approximately 100–150 larvae were put on each vole. For the first infestation, voles were anesthetized with the ketamine/xylazine cocktail described previously. Unexpectedly, of the remaining 34 voles, another 6 died under anesthesia leaving 28 animals. For the second and third infestations, voles were anesthetized with 2% isoflurane using the Combi-vet® anesthesia system (Rothacher Medical, Switzerland) and no additional animals were lost. For each vole, an ear biopsy and a blood sample (from the saphenous vein) were collected on four occasions at −1, 26, 51, and 106 days PI. Ear tissue biopsies and serum samples were kept at –20°C for future analysis. On day 106 PI, the voles were euthanized with CO_2_ and exsanguinated. Of the 28 voles that survived to the end of the study, there were 14 in the uninfected control group and 14 in the *B. afzelii*-infected group.

### Tick phenotypes

Engorged larval ticks detached from voles after 2–4 days of blood feeding. For each of the 84 combinations of infestation and vole, a maximum of 50 engorged larvae were collected and placed in individual 1.5 ml Eppendorf tubes. The tubes contained a strip of moistened paper towel to maintain a high humidity. The engorged larvae were kept in an incubator (Sanyo, Japan) with a day-night cycle of 16 hours of light and 8 hours of darkness, and a relative humidity of 85% (section 1 in the supplementary material). For each infestation and each vole, a maximum of 20 engorged larvae were randomly selected and weighed within 5 days of drop off (total of 1680 engorged larvae). Once the engorged larval ticks started molting, they were checked two or three times per week until the percentage of molted nymphs reached 80%. The median molting time was defined as the date when 50% of the engorged larvae had molted into nymphs. At two weeks after the median molting time, the ticks that had been weighed as larvae were weighed again as nymphs. At four weeks after the median molting time, 10 nymphs were randomly selected for each of the 84 combinations of vole and infestation (total of 840 nymphs), and these nymphs were frozen at –80°C to assess their *B. afzelii* infection status.

Different tick phenotypes were used to determine the effect of acquired anti-tick immunity and *B. afzelii* infection on tick fitness. The tick phenotypes included: engorged larval weight, unfed nymphal weight, molting time and molting success. Molting time was defined as the number of days between the detachment of the engorged larvae and the molting to unfed nymphs. Molting success was the percentage of engorged larvae that molted into flat nymphs.

### ELISA and qPCR

The serum samples of the voles were tested for the presence of *B. afzelii*-specific IgG antibodies using a commercial ELISA assay as previously described (31). The DNA was extracted from the vole tissue samples, the engorged nymphs, and the flat nymphs as previously described (31, 36). All the DNA extractions were eluted into 65 μl of water. The vole ear tissue samples and the ticks were tested for infection with *B. afzelii* using a qPCR assay that targets a 132 bp fragment of the *flagellin* gene as previously described (31). For each vole tissue sample and tick, 3.0 μl of DNA template was used in the qPCR reaction.

### Culture of viable *B. afzelii* spirochetes from xenodiagnostic nymphs

To demonstrate that the voles transmitted a viable *B. afzelii* infection to the *I. ricinus* ticks, flat nymphs (that had fed as larvae on the voles) were cultured in BSK-H media (Bio&Sell). Six nymphs were randomly selected for each bank vole (total of 168 nymphs). Each tick was washed with 70% ethanol, cut in half, and placed in an individual 1.5 ml Eppendorf tube containing BSK-H medium. The culture tubes were kept at 34°C in an incubator and were screened for motile *B. afzelii* spirochetes over a period of 4 weeks using a dark field microscope.

### Tick salivary gland extract protein solution

Tick salivary gland extract (SGE) proteins were obtained from the salivary glands of engorged adult female *I. ricinus* ticks from our laboratory colony. A total of 28 female and 5 male adult ticks were fed on one rabbit. After 96 hours of blood feeding, the 24 engorged female ticks were removed from the rabbit with tweezers, washed with 70% ethanol, and washed with 1x PBS. The tick salivary glands were dissected on ice using a dissecting microscope, washed with 1x PBS, pooled and homogenized in 400 µl of 1x PBS, and kept at –80 °C. The tick salivary gland cells were lysed by keeping the suspension on ice and conducting three short pulse sonication sessions (5 s pulse, 30 s pause) using a SKAN sonifier 450 (SKAN AG, Switzerland). Soluble antigens were obtained after centrifugation at 20,000 g for 15 min. The protein concentration was determined using the Bradford assay. The concentration of the tick SGE sample was 1.62 mg/ml for a total of 648 µg of tick SGE.

### Tick salivary gland extract ELISA

The IgG antibody response of the voles against tick SGE was measured using a homemade ELISA. 96-well tissue culture plates (Fisher Scientific, Switzerland) were coated overnight at 4°C with 1 μg of tick SGE protein per well. Wells were washed 3 times with PBS-Tween 0.1% between each change of solution. The wells were incubated with a BSA 2% blocking solution for 2 hours, followed by the bank vole serum samples (diluted 1:100 in 1x PBS) for 45 minutes, and the secondary antibody for 45 minutes (diluted 1:5000 in 1x PBS). The secondary antibody was a goat anti-*Mus musculus* IgG conjugated to horseradish peroxidase (Promega). After adding 100 μl of TMB product (Promega) to each well, the absorbance was measured at 652 nm every 2 minutes for one hour using a plate reader (Synergy HT, Multi-detection plate reader, Bio-Tek, United States). The strength of the IgG antibody response against tick SGE was determined for each serum sample by calculating the area under the curve of the optical density (OD) versus time.

### Infection status of bank voles

A vole was considered to be infected with *B. afzelii*, if it tested positive for one or more of four criteria: (1) optical density > 500 absorbance units indicating the presence of *B. afzelii*-specific IgG antibodies, (2) spirochete load in the ear tissue biopsy > 0, (3) spirochete load in the xenodiagnostic nymphs > 0, and (4) culture of live spirochetes from xenodiagnostic nymphs. Thirteen of the 14 voles in the *B. afzelii*-infected group tested positive for 3 or 4 criteria, and these 13 voles were therefore considered as infected with *B. afzelii* (section 2 in the supplementary material). The vole that tested negative for all five criteria was excluded from the analysis. As expected, all of the 14 voles in the control group tested negative for the 4 infection criteria. All statistical analyses are therefore based on 13 *B. afzelii*-infected voles and 14 uninfected voles.

### Statistical Analysis

All statistical analyses were done in R version 1.0.143 (R Development Core Team 2015-08-14). Means are reported with their 95% confidence intervals (95% CI). Normal versus binomial response variables were modelled using linear mixed effects models (LMMs) with normal errors versus generalized linear mixed effects models (GLMMs) with binomial errors, respectively. To determine the statistical significance of each fixed factor, pairs of models that differed in the fixed factor of interest were compared using log-likelihood ratio (LLR) tests. To calculate the p-value, the change in deviance between models is compared to the Chi-square distribution. Bank vole identity was included as a random factor.

### Tick life history traits

The response variables included four tick life history traits: (1) engorged larval weight, (2) unfed nymphal weight, (3) molting duration, and (4) molting success. The engorged larval weight and unfed nymphal weight were log10-transformed to improve the normality of the residuals. All response variables followed a normal distribution, except molting success, which followed a binomial distribution. Each response variable was modelled as a function of two fixed factors: infection status (two levels: uninfected control and infected with *B. afzelii*), infestation number (two levels: first and third infestation), and their interaction. For infestation number, only the first and third infestations were used in the analysis to simplify the presentation of the results (full analyses in section 4 of the supplementary material). We also modelled the tick SGE-specific IgG antibody levels in the voles (log10-transformed) using an LMM model. The fixed factors were infection status, day of blood sample (four levels: −1, 26, 51, and 106 days PI), and their interaction. The background variation in the OD values was corrected for each plate as follows: the mean OD of the BSA controls for a given plate was subtracted from the OD values of the bank vole serum samples in that plate.

### Relationships between *B. afzelii* infection phenotypes in the infected voles

For the subset of infected voles, we had measured three *B. afzelii* infection phenotypes around the time of each larval infestation (27, 54, and 84 days PI): (1) tick SGE-specific IgG antibody levels of the vole (hereafter the anti-tick IgG response), (2) spirochete load in the vole ear biopsy (number of spirochetes in the entire ear tissue biopsy), and (3) host-to-tick transmission (percentage of flat nymphs that were infected with *B. afzelii*). The spirochete loads in the ear biopsies were log10-transformed to improve their fit to a normal distribution. The anti-tick IgG response and the log10(ear tissue spirochete load) followed a normal distribution, whereas host-to-tick transmission followed a binomial distribution. We conducted two separate analyses that were based on causal relationships between these three variables. First, we modelled the ear biopsy spirochete load as a function of the anti-tick IgG response. Second, we modelled host-to-tick transmission as a function of the anti-tick IgG response and the ear biopsy spirochete load.

## Results

### Collection of engorged nymphal ticks from the voles

We collected 47 engorged nymphs from the 14 voles in the control group (mean = 3.36, range = 1–5 engorged nymphs per vole). As expected, none of these nymphs tested positive for *B. afzelii*. We collected 46 engorged nymphs from the 14 voles in the *B. afzelii*-infected group (mean = 3.29, range = 0–5 engorged nymphs per vole). Our qPCR assay found that 58.7% (27/46) of the engorged nymphs were infected with *B. afzelii*. For 12 of the 14 voles in the infected group, we collected at least 1 engorged *B. afzelii*-infected nymph (mean = 1.93, range = 1–4; Table S1 in the supplementary material). These data show that the voles in the infected group and the control group were infested with similar numbers of nymphs.

### Life history traits of *I. ricinus* ticks that fed on bank voles

#### Weight of blood-engorged larvae

The engorged larval weight decreased from the first to the third infestation (Figure 1). The LMM analysis of engorged larval weight found a significant interaction between infection status and infestation number (LLR: χ^2^ = 8.316, df = 2, p = 0.016). Infestation number had a significant effect on the engorged larval weight (LLR: χ^2^ = 45.996, df = 2, p < 0.001), but the effect of infection status was not significant (LLR: χ^2^ = 0.028, df = 1, p = 0.866). For the first, second, and third larval infestation, the mean engorged larval weight was 483 μg (95% CI: 475–492), 452 μg (95% CI: 444–461), and 456 μg (95% CI: 448–465), respectively. Compared to the first infestation, the mean engorged larval weight in the second and third infestation was reduced by 6.4% (p = 0.006) and 5.6% (p < 0.001), respectively.

**Figure 1.**
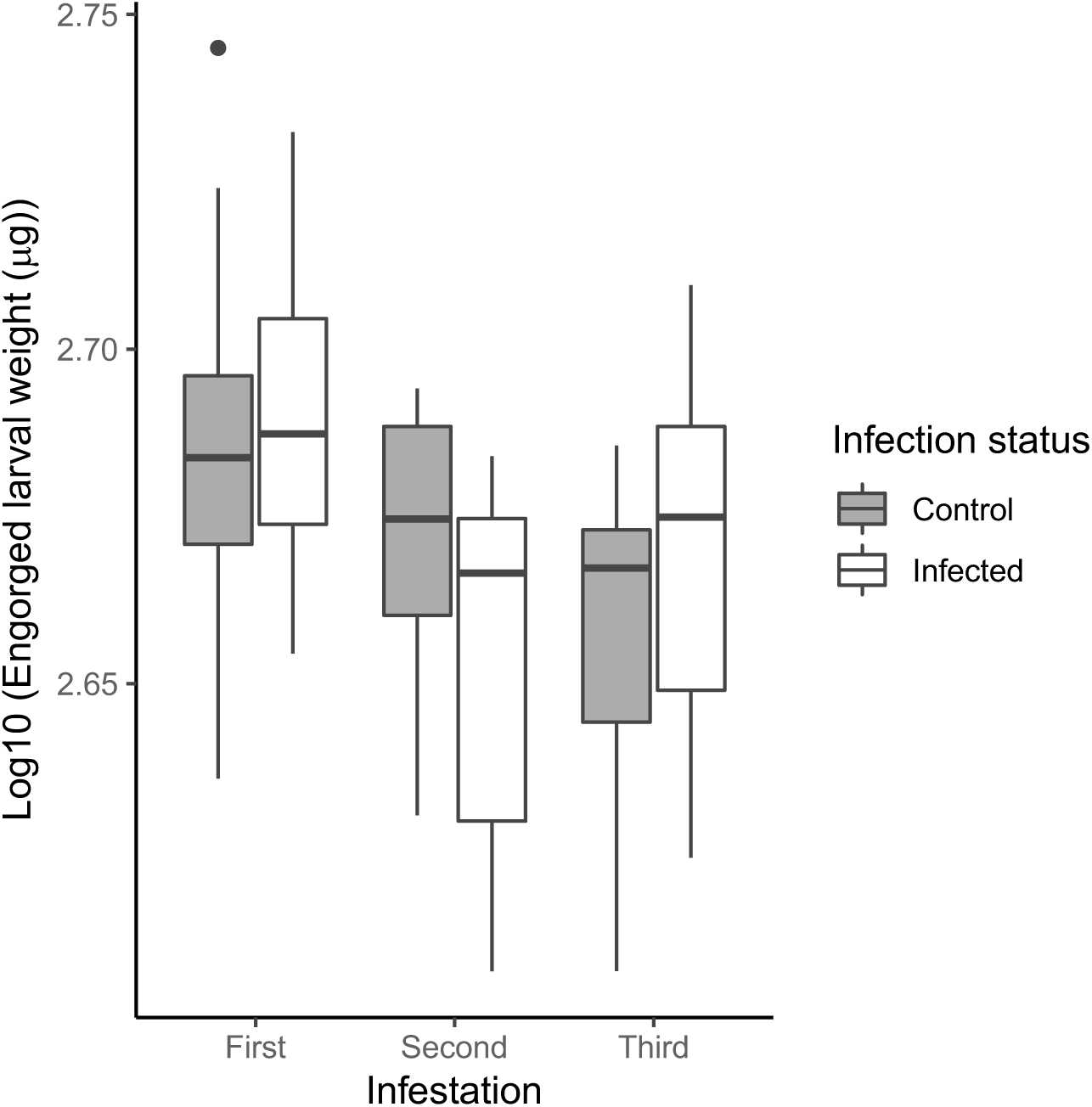
The engorged weight of *I. ricinus* larval ticks feeding on the bank voles decreased over the three successive infestations. Infection of bank voles with *B. afzelii* did not affect the engorged larval weight (measured in μg). The bank voles in the *B. afzelii*-infected group (n = 13) and the control group (n = 14) were infested with larval ticks at 27, 55, and 84 days PI. Each data point represents the mean engorged larval weight for an individual bank vole and is based on ∼20 ticks. Shown are the medians (black line), the 25th and 75th percentiles (edges of the box), the minimum and maximum values (whiskers), and the outliers (circles).

#### Weight of the unfed nymphs

The flat nymphal weight decreased from the first to the third infestation (Figure 2). The LMM analysis of flat nymphal weight found no significant interaction between infection status and infestation number (LRR: χ^2^ = 5.299, df = 2, p = 0.071). After removing the interaction, infestation number had a significant effect on the flat nymphal weight (LRR: χ^2^ = 74.807, df = 2, p < 0.001), but the effect of infection status was not significant (LRR: χ^2^ = 0.656, df = 1, p = 0.418). For the first, second, and third larval infestation, the mean nymphal weight was 188 μg (95% CI: 183– 192), 166 μg (95% CI: 162–171), and 177 μg (95% CI: 172–181), respectively. Compared to the first infestation, the mean flat nymphal weight in the second and third infestation was reduced by 11.7% (p < 0.001) and 5.9% (p < 0.001), respectively.

**Figure 2.**
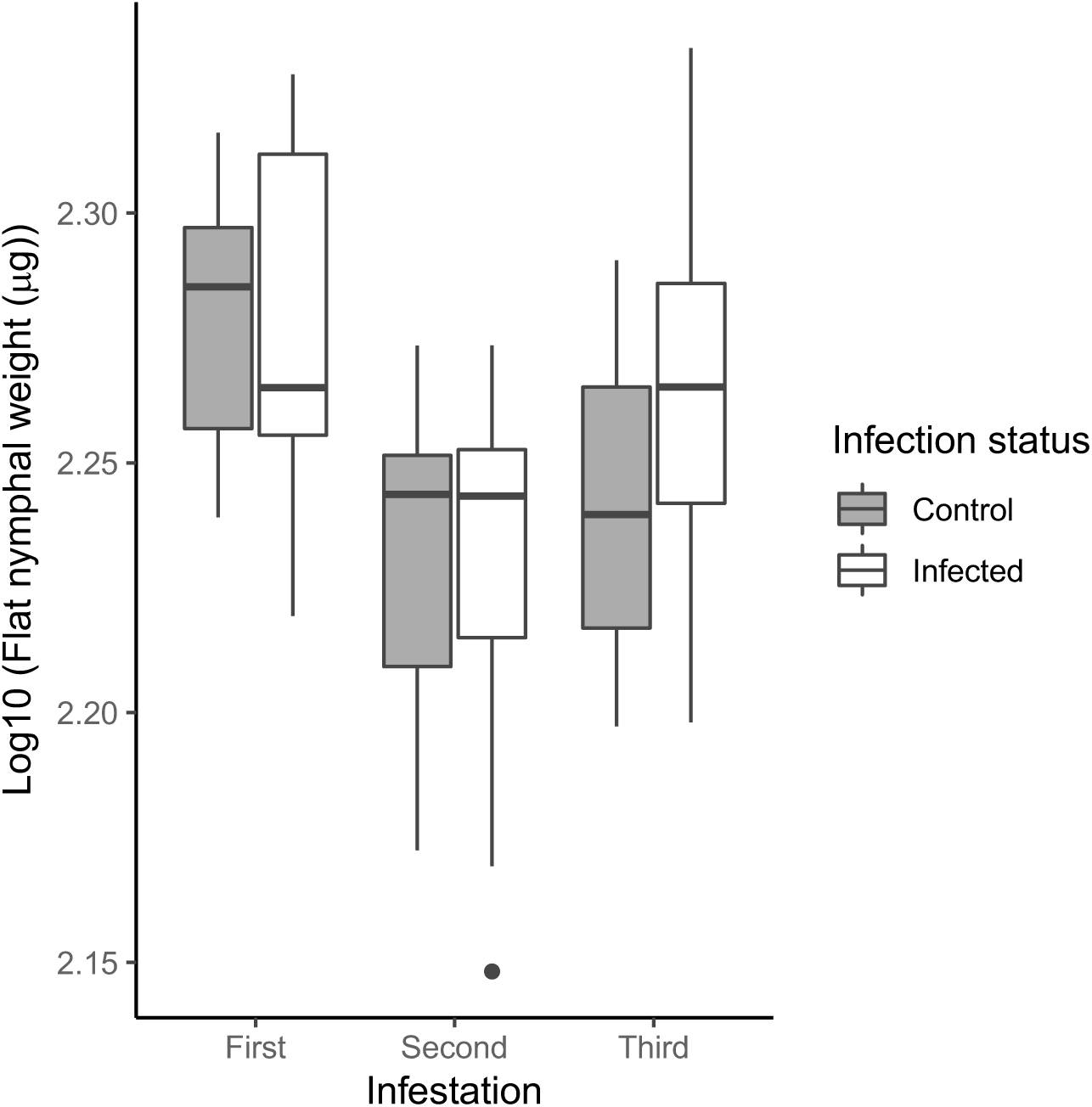
The weight of the flat *I. ricinus* nymphs decreased over the three successive infestations. The flat nymphs had fed on the bank voles during the larval stage. Infection of bank voles with *B. afzelii* did not affect the flat nymphal weight (measured in μg). The bank voles in the *B. afzelii*-infected group (n = 13) and the control group (n = 14) were infested with larval ticks at 27, 55, and 84 days PI. Each data point represents the mean flat nymphal weight for an individual bank vole and is based on ∼20 flat nymphs. Shown are the medians (black line), the 25th and 75th percentiles (edges of the box), the minimum and maximum values (whiskers), and the outliers (circles).

#### Molting time of engorged larval ticks to nymphs

The larva-to-nymph molting time was defined as the number of days between the drop-off of the engorged larval ticks and the molt into flat nymphs. The molting time was monitored for a total of 2611 engorged larval ticks, of which 83.80% (2188/2611) molted into nymphs. The molting time decreased over the three successive larval infestations (Figure 3). The LMM analysis of molting time found no significant interaction between infection status and infestation number (LRR: χ^2^ = 4.341, df = 2, p = 0.114). After removing the interaction, infestation number had a significant effect on the molting time (LRR: χ^2^ = 246.7, df = 2, p < 0.001), but the effect of infection status was not significant (LRR: χ^2^ = 0.004, df = 1, p = 0.946). For the first, second, and third larval infestation, the mean molting time was 51 days (95% CI: 50–53), 47 days (95% CI: 45–49), and 35 days (95% CI: 33–37), respectively. Compared to the first infestation, the mean molting time in the second and third infestation was reduced by 7.8% (p < 0.001) and 31.4% (p < 0.001), respectively.

**Figure 3.**
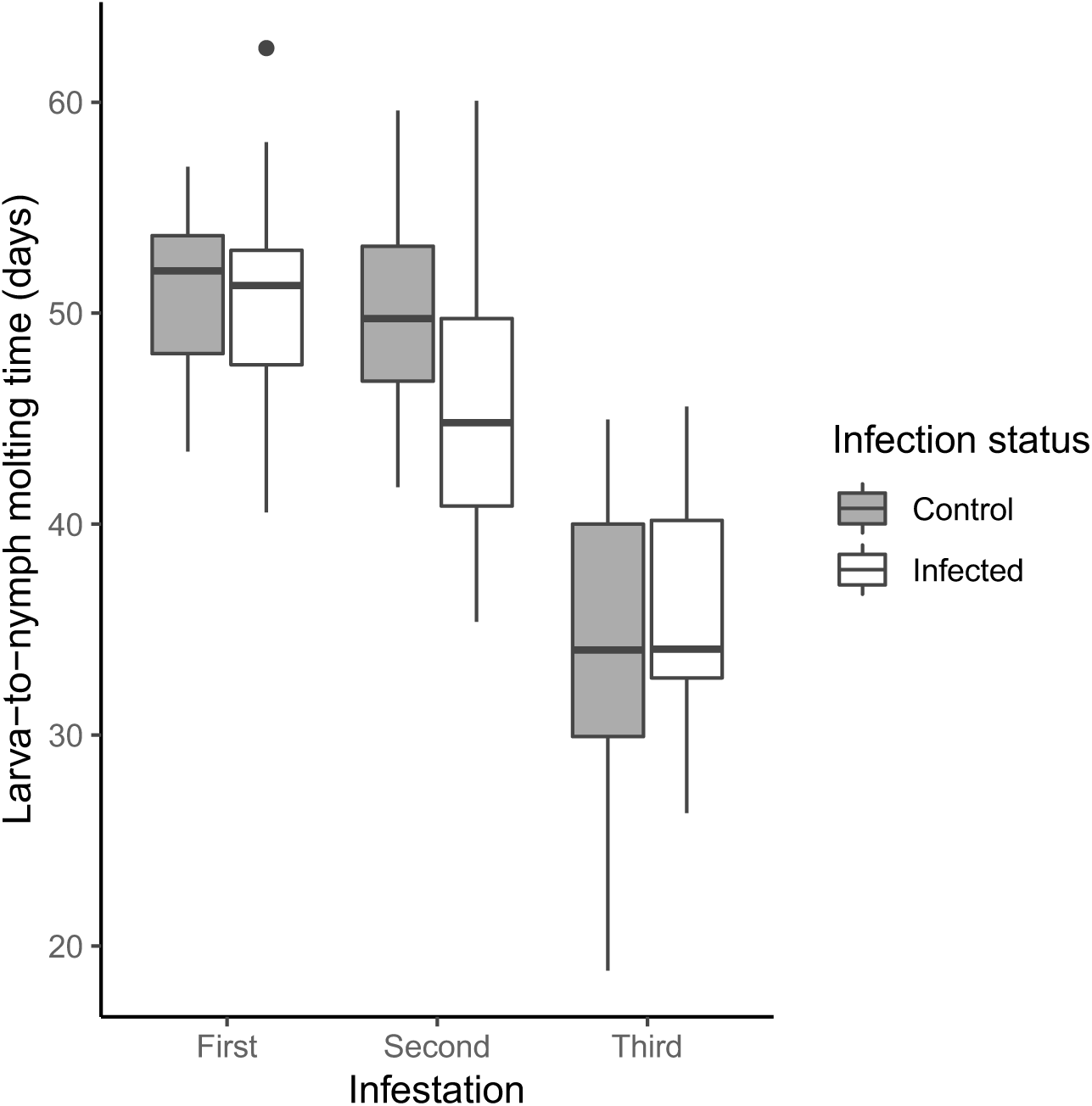
The larva-to-nymph molting time of *I. ricinus* ticks decreased over the three successive infestations. The molting time refers to the number of days for an engorged larval tick to molt into a nymphal tick. Infection of bank voles with *B. afzelii* did not affect the molting time. The bank voles in the *B. afzelii*-infected group (n = 13) and the control group (n = 14) were infested with larval ticks at 27, 55, and 84 days PI. Each data point represents the mean larva-to-nymph molting time for an individual bank vole and is based on ∼50–100 ticks. Shown are the medians (black line), the 25th and 75th percentiles (edges of the box), the minimum and maximum values (whiskers), and the outliers (circles).

#### Larva-to-nymph molting success

The molting success was defined as the percentage of engorged larval ticks that molted into flat nymphs. The molting success decreased over the three successive larval infestations (Figure 4). The GLMM analysis of molting success found no significant interaction between infection status and infestation number (LRR: χ^2^ = 2.276, df = 2, p = 0.320). After removing the interaction, infestation number had a significant effect on the molting success (LRR: χ^2^ = 35.121, df = 2, p < 0.001), but the effect of infection status was not significant (LRR: χ^2^ = 1.489, df = 1, p = 0.222). For the first, second, and third larval infestation, the mean molting success was 88% (95% CI: 86–90%), 87% (95% CI: 84–91%), and 78% (95% CI: 75–81%), respectively. Compared to the first infestation, the mean molting success in the second and third infestation was reduced by 1.1% (p = 0.883) and 11.4% (p < 0.001), respectively.

**Figure 4.**
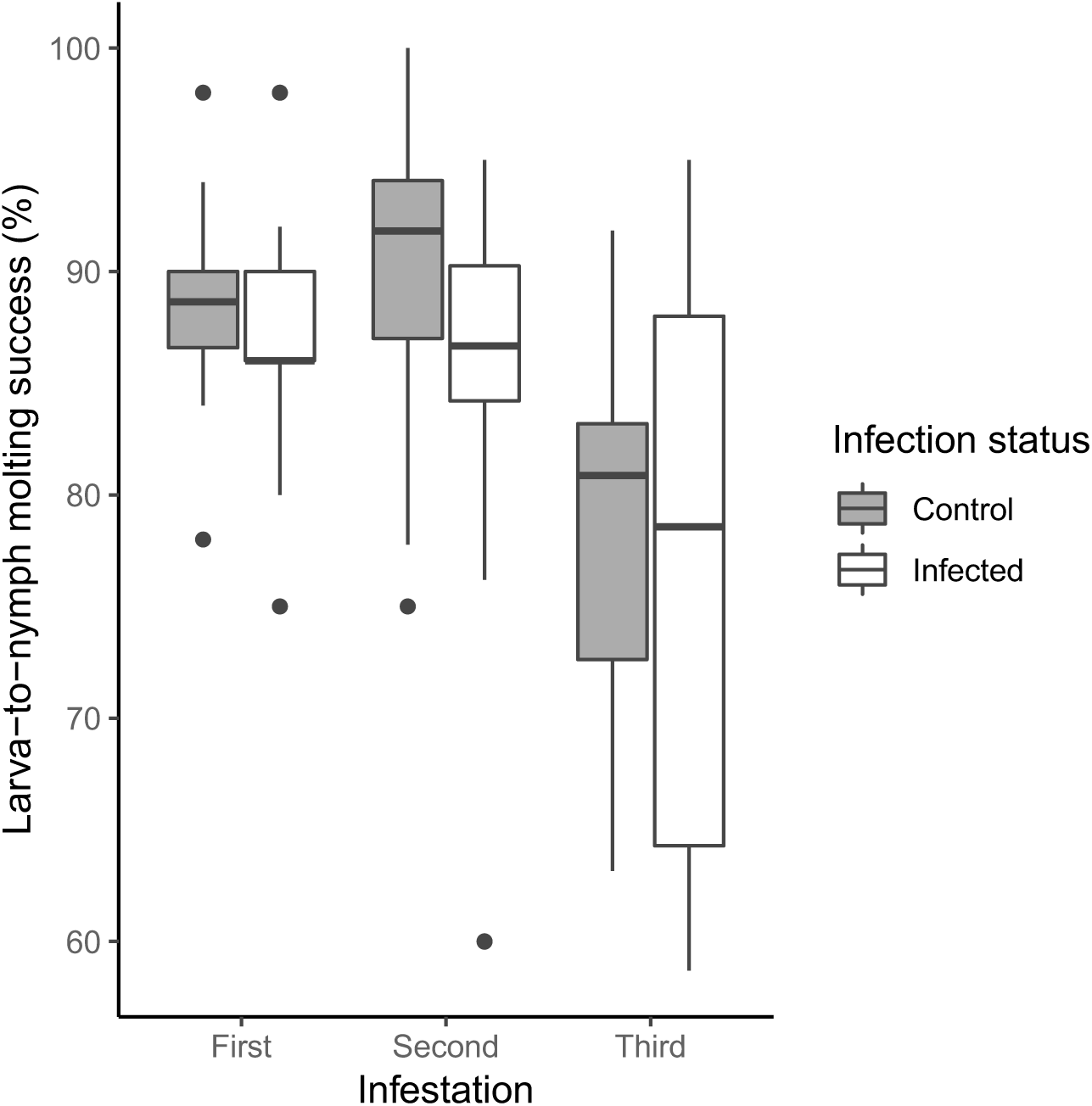
The molting success of *I. ricinus* larval ticks decreased over the three successive infestations. Molting success refers to the percentage of engorged larval ticks that developed into the nymphal stage. Infection of bank voles with *B. afzelii* did not affect the molting success. The bank voles in the *B. afzelii*-infected group (n = 13) and the control group (n = 14) were infested with larval ticks at 27, 55, and 84 days PI. Each data point represents the mean larva-to-nymph molting success for an individual bank vole and is based on ∼50–100 ticks. Shown are the medians (black line), the 25th and 75th percentiles (edges of the box), the minimum and maximum values (whiskers), and the outliers (circles).

#### Bank voles developed a strong IgG antibody response against the salivary gland extract of *I. ricinus* ticks

The tick SGE-specific IgG antibody levels (hereafter the anti-tick IgG response) increased over the duration of the study (from day −1 to day 106; Figure 5) and this change was significant (LME: χ^2^ = 166.61, df = 1, p < 0.001). For day −1, day 26, day 51, and day 106, the mean anti-tick IgG response (measured in absorbance units) was 1176 (95% CI: 1040–1328), 1559 (95% CI: 1370–1774), 2391 (95% CI: 2112–2706), and 6351 (95% CI: 5635–7157), respectively. Compared to day - 1, the mean anti-tick IgG response on day 106 had increased 5.4-fold.

**Figure 5.**
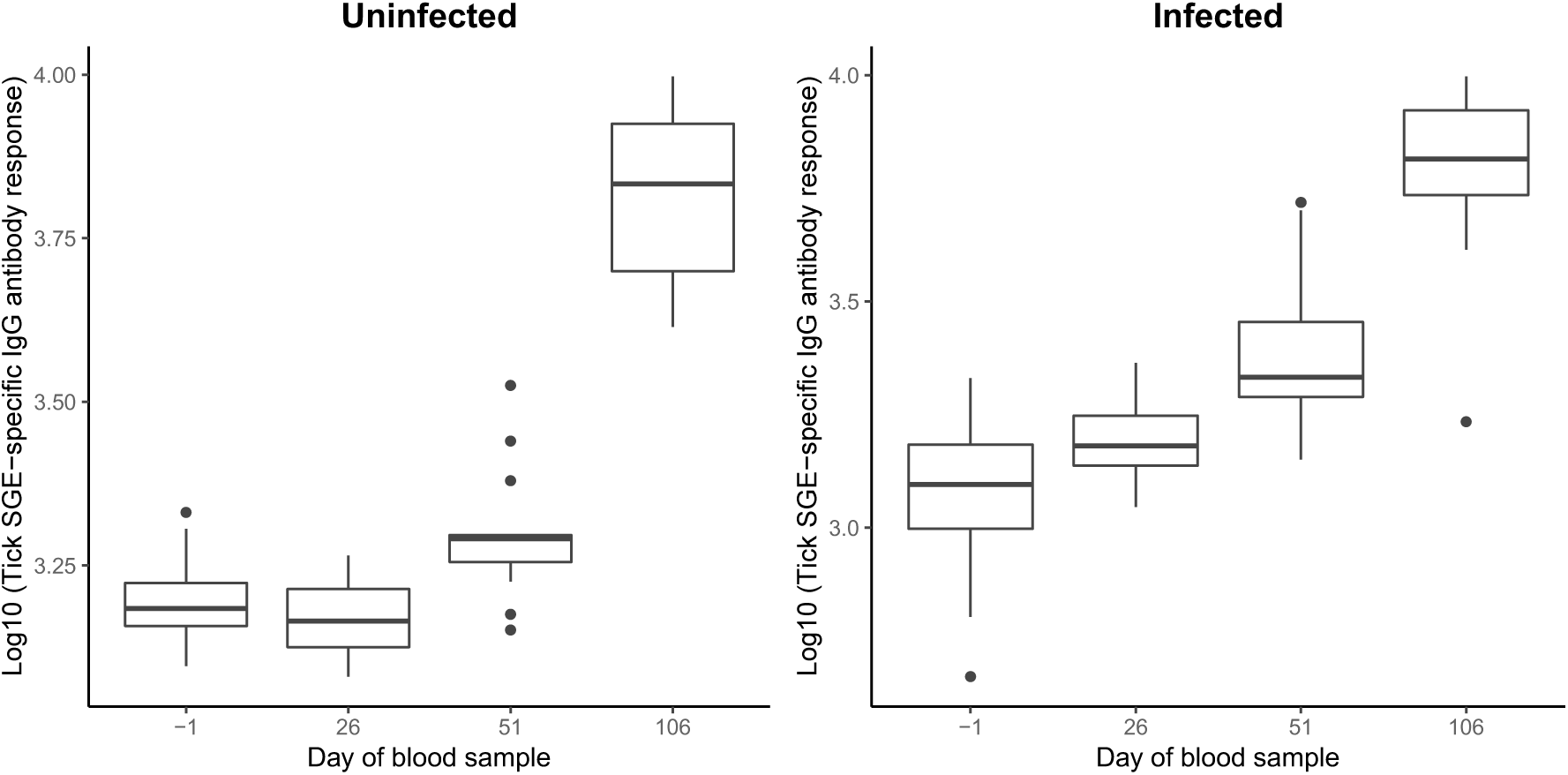
The bank voles developed a strong IgG antibody response against the salivary gland extract of *I. ricinus* ticks over the three successive larval infestations in both the control group and the *B. afzelii*-infected group. Serum samples were taken on −1, 26, 51, and 106 days PI, which corresponded to time points before the nymphal challenge, and after the first, second, and third larval infestation. The strength of the IgG antibody response against tick salivary gland extract was measured as the absorbance from an ELISA. Each data point represents the log10-transformed absorbance value for an individual bank vole. Shown are the medians (black line), the 25th and 75th percentiles (edges of the box), the minimum and maximum values (whiskers), and the outliers (circles).

The LMM analysis of the anti-tick IgG response found that the interaction between infection status and the day of the blood sample was significant (LRR: χ^2^ = 12.638, df = 3, p = 0.005). We therefore tested the effect of infection status on the anti-tick IgG response separately for each day. This analysis found a significant difference between the infected group and the control group at day −1 (t-test: df = 25, t = 6.818, p < 0.001), day 26 (t-test: df = 22, t = 2.218, p = 0.037), day 51 (t-test: df = 24, t = 3.1036, p = 0.004), but not at day 106 (t-test: df = 25, t = 0.117, p = 0.908). Thus, the anti-tick IgG response reached similar levels in both groups of bank voles at the end of the experiment. On day 26 and day 51, the mean anti-tick IgG response of the infected group was 17.4% and 45.6% higher than the control group, respectively (Figure 5). On day −1, the mean anti-tick IgG response of the infected group was 55.6% lower than the control group (Figure 5).

### Abundance of *B. afzelii* in ear biopsies and host-to-tick transmission

#### Spirochete load in the bank vole ear biopsy

The spirochete loads in the vole ear biopsies (2 mm diameter) decreased over the three infestations (section 5 in the supplementary material). For the first, second, and third larval infestation, the mean spirochete load in the ear biopsy (measured in number of spirochetes) was 5883 (95% CI: 1516–22834), 202 (95% CI: 52–786), and 13 (95% CI: 3–50), respectively. The anti-tick IgG response had a significant negative effect on the spirochete load in the ear biopsy (Figure 6; LME: χ^2^ = 17.748, df = 1, p < 0.001; slope ± S.E. = −2.26 ± 0.537).

**Figure 6.**
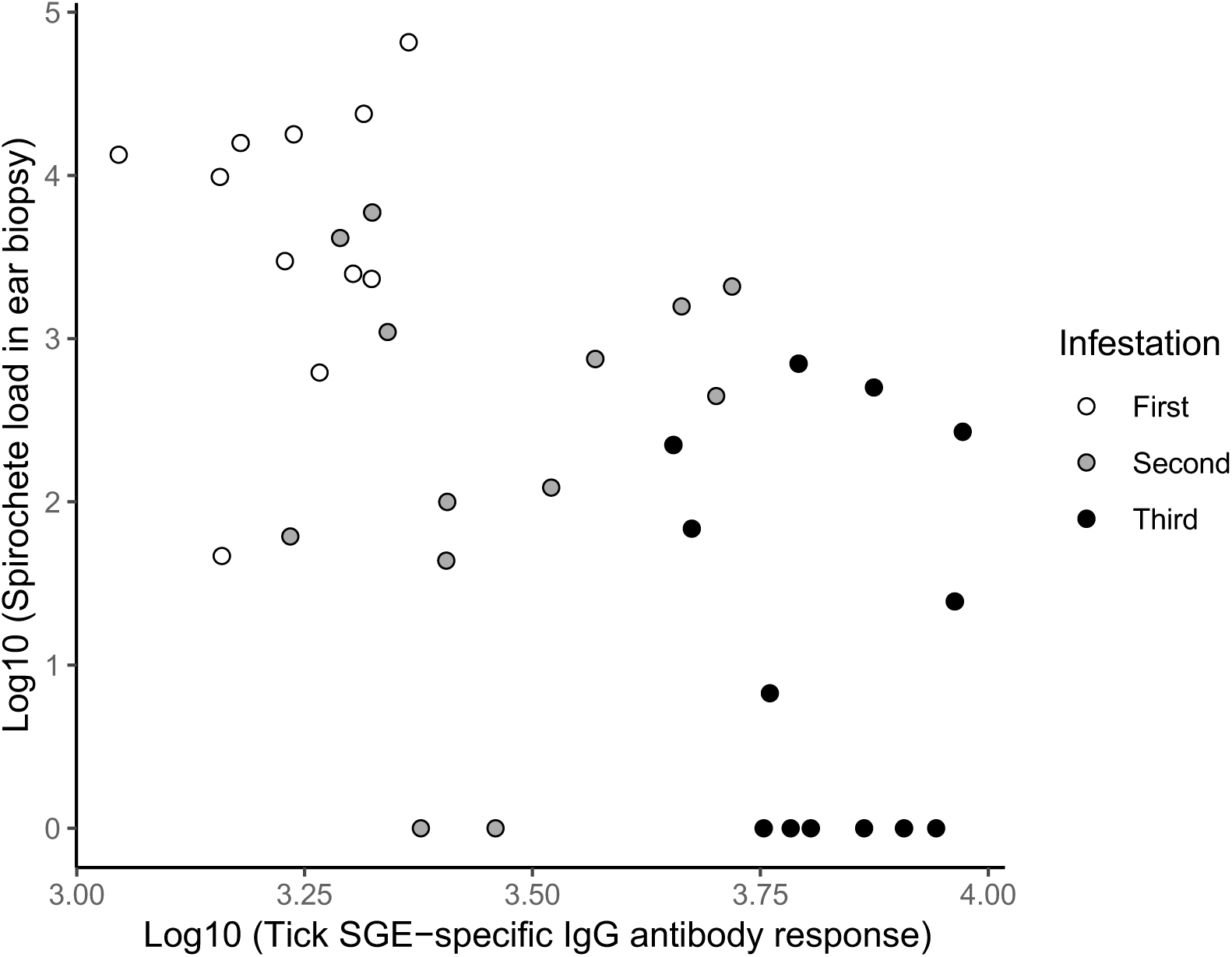
The spirochete load in the bank vole ear biopsy is negatively related to the tick SGE-specific IgG antibody response. Data are shown for the subset of *B. afzelii*-infected bank voles (n = 13 individuals) at the time of the first (open white circles), second (solid grey circles), and third infestation (solid black circles). Ear tissue biopsies were taken at 26, 51, and 106 days PI and the serum samples were taken at 26, 51, and 106 days PI. The 39 data points represent the 13 *B. afzelii*-infected bank voles at each of the three infestations. Spirochete loads refer to the number of spirochetes in the whole ear tissue biopsy (2 mm diameter).

#### Host-to-tick transmission of *Borrelia afzelii*

After combining the three infestations, host-to-tick transmission was 38.5% (144 infected nymphs/ 374 total nymphs). Host-to-tick transmission decreased over the three infestations (section 6 in the supplementary material). For the first, second, and third larval infestations, the host-to-tick transmission was 59.4% (76/128), 35.6% (42/118), and 20.3% (26/128), respectively. For 30.8% (4/13) of the voles, the host-to-tick transmission was 0% by the third infestation. The anti-tick IgG response had a significant negative effect (GLME: χ^2^ = 5.407, df = 1, p = 0.020; logit slope ± S.E. = −1.49 ± 0.639), whereas the log10-transformed spirochete load in the ear biopsy had a significant positive effect on host-to-tick transmission (Figure 7; GLME: χ^2^ = 10.369, df = 1, p = 0.001; logit slope ± S.E. = 0.52 ± 0.162).

**Figure 7.**
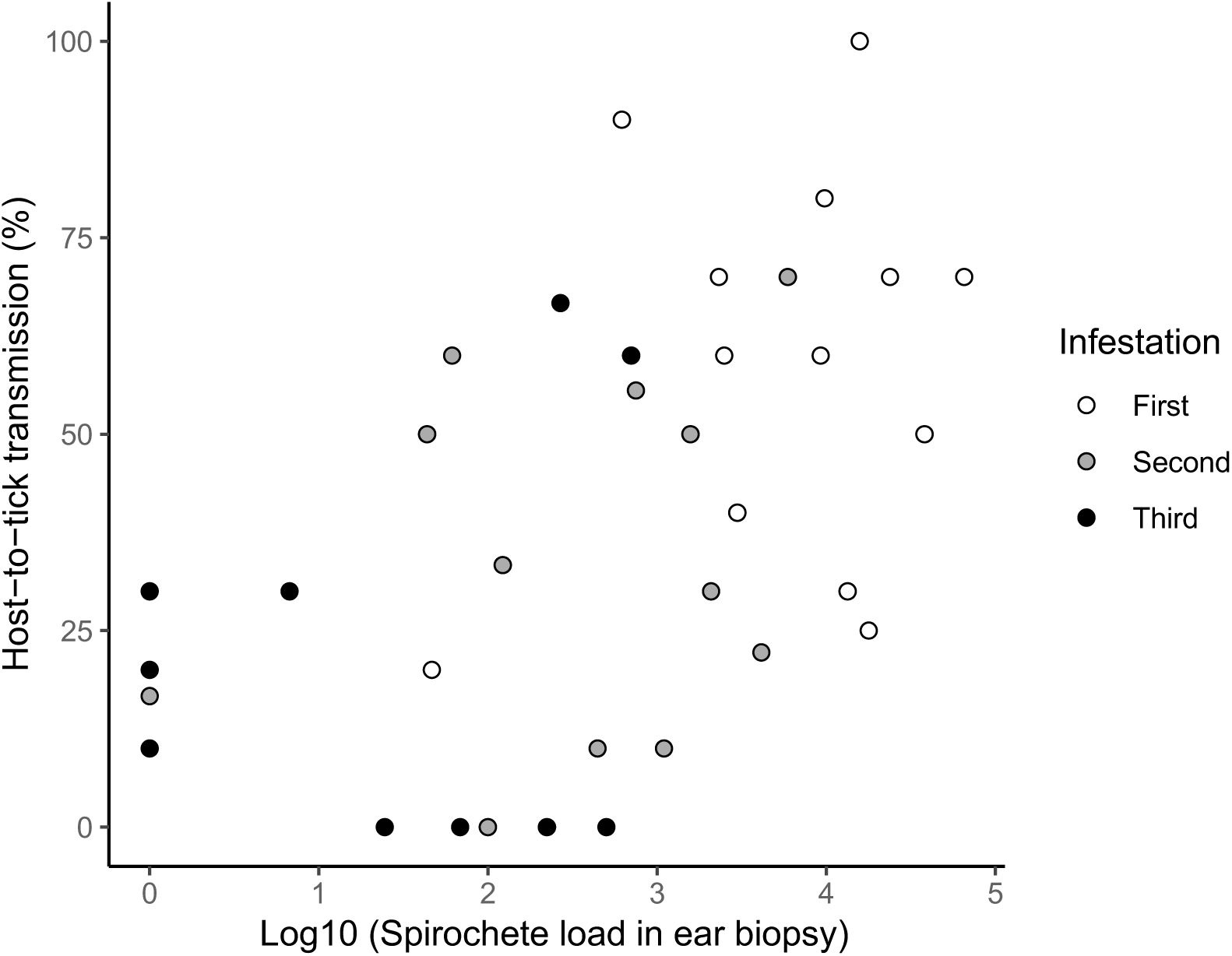
Host-to-tick transmission is positively correlated with spirochete load in the ear tissue of the bank voles. Host-to-tick transmission is the percentage of nymphs that acquired the *B. afzelii* infection during the larval blood meal. Data are shown for the subset of *B. afzelii*-infected bank voles (n = 13 individuals) at the time of the first (open white circles), second (solid grey circles), and third infestation (solid black circles). Ear tissue biopsies were taken at 26, 51, and 106 days PI and the bank voles were infested with larval ticks at 27, 55, and 84 days PI. The 39 data points represent the 13 *B. afzelii*-infected bank voles at each of the three infestations. Spirochete loads refer to the number of spirochetes in the whole ear tissue biopsy (2 mm diameter).

## Discussion

### Ability of *B. burgdorferi* sl to suppress acquired immunity in the vertebrate host

Infection with *B. afzelii* in a natural rodent host did not prevent the development of acquired immunity against larval *I. ricinus* ticks and its negative effects on tick life history traits. Our study contrasts with a recent study showing that *B. burgdorferi* ss suppressed the development of acquired immunity in laboratory *Mus musculus* mice (26). Specifically, this study found that infected mice immunized with the influenza vaccine did not develop protective antibodies against the influenza virus (26). The mechanism of immunosuppression is that *B. burgdorferi* ss migrates to the mouse lymph nodes where it inhibits the development of long-lived plasma cells and memory B cells, (26). Differences between our study and (26) include the genospecies of *B. burgdorferi* sl (*B. burgdorferi* ss versus *B. afzelii*), the antigen (influenza vaccine versus live ticks), the rodent host (lab mice versus natural host), and mode of infection (needle inoculation versus tick bite). Other tick-borne pathogens such as *Anaplasma phagocytophilum* and *Babesia microti* have been documented to induce immunosuppression in vertebrate hosts (23, 37). Interestingly, the protozoan parasite *B. microti* was able to induce immunosuppression in laboratory mice, but not in the bank vole or any known natural reservoir host (37, 38). These studies suggest that the ability of *B. burgdorferi* sl to suppress the adaptive immune system may depend on the identity of the vertebrate host.

From an ecological and public health perspective, pathogen-induced immunosuppression is important because it can facilitate the infection and emergence of opportunistic pathogens (39). For example, the ability of HIV to suppress the immune system has led to the re-emergence of tuberculosis in human populations in the developing world (40). Community ecology studies often find positive associations between *B. burgdorferi* sl genospecies and other tick-borne pathogens (39). For example, the emergence of *B. microti* was strongly associated with the prevalence of *B. burgdorferi* ss in the northeast United States (41, 42). In addition, *B. burgdorferi* sl genospecies and strains within a genospecies have been found to be positively associated (43-45). In summary, *B. burgdorferi* sl-induced immunosuppression in the vertebrate host can facilitate mixed infections and the emergence of other tick-borne pathogens.

### Development of acquired immunity against ticks in vertebrate hosts

Our study found that repeated infestations with larval *I. ricinus* ticks caused the bank voles to develop a strong IgG antibody response against the SGE proteins of *I. ricinus*. The development of this anti-tick immunity was correlated with a reduction in tick fitness including the size of the engorged larval ticks and the resultant flat nymphs, the duration of the larva-to-nymph molt, and the larvae-to-nymph molting success. One complication with our study was that the *I. ricinus* larvae for the first and second infestation came from the University of Neuchâtel colony and the larvae for the third infestation came from Insect Services (Germany). This happened because the University of Neuchâtel colony failed to produce sufficient numbers of larvae for the third infestation. As a result, any differences in tick phenotype between the first and third infestation could be due to innate differences between these two tick colonies rather than acquired anti-tick immunity. However, the statistically significant decreases in engorged larval weight and flat nymphal weight between the first and second larval infestation (which used larvae from the University of Neuchâtel colony) suggests that our bank voles developed acquired anti-tick immunity after the first larval infestation.

Our results are in agreement with previous studies showing that bank voles develop anti-tick immunity to larval ticks of *I. ricinus* or *I. trianguliceps* (14, 27, 37). These studies found that anti-tick immunity reduced the percentage of fully engorged larvae, larval engorgement index, duration of attachment, percentage of recovered larvae, and larva-to-nymph molting success (14, 27, 37). The mechanism by which anti-tick immunity reduces tick fitness is that it impairs tick blood feeding and therefore reduces the quality and quantity of the blood meal (21). With respect to body size, previous studies on *I. ricinus* have shown that larger nymphs have higher fat reserves (46) and that such nymphs can quest for longer periods of time, which increases their chances of finding a host in the field (9, 47, 48). Body size also influences female fecundity, with larger female ticks laying larger clutches of eggs (49). The reduced molting success is the most direct fitness cost because for arthropods such as ticks, the inability to complete the molt is equivalent to death (50). Population ecology models of *Ixodes* ticks have shown that the tick population growth rate is highly sensitive to molting success (51, 52). In summary, the negative effects of acquired anti-tick immunity on tick life history traits can reduce the abundance of ticks in the field.

### *B. burgdorferi* sl pathogens establish chronic infections in rodents with high lifetime host-to-tick transmission

We showed that *B. afzelii* isolate NE4049 established a long-lived systemic infection in the bank voles. At the time of sacrifice, all of the 13 bank voles showed systemic infections where the dorsal skin, ventral skin, ear, bladder and heart were all infected with *B. afzelii*. These results are in agreement with previous studies that have shown that *B. burgdorferi* sl pathogens established chronic systemic infections in their rodent reservoir hosts (27, 33, 53, 54). For vector-borne pathogens such as *B. burgdorferi* sl, chronic infections are adaptive because they increase the lifetime transmission success of the infection (24, 33, 55). In the present study, host-to-tick transmission of *B. afzelii* decreased three-fold over time from 60% in the first infestation (4 weeks PI) to 20% in the third infestation (12 weeks PI). Less dramatic declines have been observed in another important rodent reservoir, the wood mouse (*Apodemus sylvaticus*), where host-to-tick transmission of *B. afzelii* decreased from 100% at 3 weeks PI to ∼40% at 9 weeks PI (54). A study that used the same isolate of *B. afzelii* (NE4049) and the same colony of *I. ricinus* ticks to infect *M. musculus* mice found that host-to-tick transmission decreased from 90.8% at 5 weeks PI to 68.9% at 13 weeks PI (33). Likewise, experimental infection studies on *B. burgdorferi* ss have shown that the pattern of host-to-tick transmission over time can vary depending on the particular combination of strain and rodent species (56-58). In general, host-to-tick transmission of *B. burgdorferi* sl pathogens decreases over time in most rodent species, but the bank voles in this study showed a particularly steep decline.

### Relationship between pathogen abundance in host tissues and host-to-vector transmission

For vector-borne pathogens, there is a direct relationship between the pathogen’s abundance in the relevant tissues and host-to-vector transmission. For example, this relationship has been shown in rodent malaria where the parasite density in the blood is critical for mouse-to-mosquito transmission (59). For *B. burgdorferi* sl pathogens, the skin rather than the blood is the critical tissue for host-to-tick transmission (60, 61). Field studies on *B. afzelii* in bank voles and other wild rodents found a positive relationship between the spirochete load in the ear tissues and transmission to larval *I. ricinus* ticks (62). Similarly, an infection experiment with laboratory *M. musculus* mice and the *B. afzelii* strain used in the present study (NE4049), also found a positive relationship between the spirochete load in the ears and the host-to-tick transmission (36). Studies on laboratory *M. musculus* mice have shown that the spirochete load in the host tissues can change over time (63-65), but these studies have not investigated host-to-tick transmission. The present study found that the spirochete load of *B. afzelii* in the ear biopsies decreased 450-fold over time from 5883 spirochetes per biopsy in the first infestation (4 weeks PI) to 13 spirochetes per biopsy in the third infestation (12 weeks PI). This decline in spirochete load was strongly correlated with the temporal increase of anti-tick immunity. Our study suggests that an increasingly effective host immune response (against the tick and/or the pathogen) reduced the spirochete load in the skin of the bank voles, which subsequently reduced transmission of *B. afzelii* to feeding larval ticks.

### Acquired immunity against *B. burgdorferi* sl and *Ixodes* ticks

Rodents develop strong antibody responses against *B. burgdorferi* sl pathogens, which are believed to play an important role in controlling the infection (66-68). For example, SCID mice that cannot produce antibodies have spirochete loads that are an order of magnitude higher than immunocompetent mice, and inoculation of SCID mice with *B. burgdorferi* sl anti-serum reduces the tissue spirochete load (64, 69-71). Previous studies on bank voles and *Apodemus* mice suggested that the strength of the spirochete-reactive antibody response was important for controlling host-to-tick transmission (29). The bank voles in this study developed a strong IgG antibody response against *B. afzelii* as shown by the results from the commercial ELISA. Thus, one plausible explanation is that the *B. afzelii*-targeted antibody response was able to reduce the spirochete load in the skin and thereby reduce host-to-tick transmission. A second explanation is that the anti-tick immunity developed against the sequential larval infestations reduced host-to-tick transmission success of *B. afzelii*. Previous studies have shown that the development of acquired immunity against ticks in the vertebrate host has important consequences for the acquisition and transmission of tick-borne pathogens (17, 18). We found a highly significant negative relationship between the anti-tick IgG antibody response and host-to-tick transmission, even after controlling for the temporal decline in spirochete load in the ear tissues. This observation suggests that anti-tick immunity caused the temporal decline in host-to-tick transmission. A plausible mechanism for this phenomenon is that the anti-tick antibodies opsonize the tick salivary gland proteins and thereby transform the feeding lesion into an immunologically hostile environment where spirochete survival and spirochete migration to the mouthparts of feeding larval ticks are compromised (72-74). A third explanation for the decline in host-to-tick transmission was that anti-tick immunity shortened the feeding time of the larval ticks, which reduced the probability of acquiring *B. afzelii*. A recent manipulative study showed that there was a positive relationship between the duration of the blood meal of *I. scapularis* larvae and the probability of host-to-tick transmission of *B. burgdorferi* ss (75). In summary, anti-spirochete immunity, anti-tick immunity, and shorter feeding times of larval ticks are three alternative explanations for the observed temporal decline in the ear tissue spirochete loads and host-to-tick transmission of *B. afzelii*.

### Ecological consequences of acquired immunity in bank voles and associated reductions in tick life history traits and host-to-tick transmission

Several studies have suggested that the bank vole is an important reservoir host of *B. afzelii* (14, 27, 29, 76-79). Early studies on the vertebrate host community in Sweden suggested that the bank vole was the second-most important host of *B. afzelii* and produced 17% of the infected *I. ricinus* nymphs (76). In this experimental study, we found that the anti-tick immunity developed by bank voles had negative consequences for both the tick vector and the tick-borne pathogen. Population ecology models of *I. ricinus* have shown that the tick population size and growth rate are highly sensitive to density-dependent mortality on the host, which can be mediated by acquired immunity (52). A recent field study in France found that the abundance of bank voles had negative effects on the abundance of *I. ricinus* nymphs the following year (80). The explanation was that acquired immunity in bank voles had a negative effect on the recruitment of feeding larval ticks into nymphs the following year. Theoretical models have shown that the reproduction number (R_0_) of *B. burgdorferi* sl pathogens is strongly influenced by the duration of the infection and the host-to-tick transmission rate (81). Both of these life history traits were strongly reduced in the present study showing the importance of acquired immunity in controlling the ecology of *B. burgdorferi* sl pathogens and the epidemiology of Lyme borreliosis.

## Conclusions

We found no evidence that infection with *B. afzelii* suppressed the development of acquired immunity against *I. ricinus* ticks in bank voles. Future research needs to confirm whether suppression of acquired immunity is a widespread strategy in the *B. burgdorferi* sl genospecies complex or whether it is restricted to certain strains or unnatural hosts, such as laboratory mice. Repeated infestations with larval *I. ricinus* ticks induced the bank voles to develop a strong IgG antibody response against the salivary gland extract proteins of *I. ricinus*. This anti-tick antibody response reduced tick fitness and was also associated with a dramatic temporal decline in the ear spirochete load and host-to-tick transmission. Our study suggests that acquired anti-tick immunity in the vertebrate host can play an important role in controlling the abundance of *B. burgdorferi* sl-infected nymphs and hence the risk of Lyme borreliosis.

## Acknowledgements

This work was supported by a grant from the Swiss National Science Foundation (SNSF) to Maarten Voordouw (FN 31003A_141153).

## Author contributions

A.G.-C. and M.J.V designed the study. A.G.-C., Y.L. and A.S performed the experimental infections and the molecular work. O.R. created the larvae necessary for the infestations. A.G.-C. analysed the data. A.G.-C. and M.J.V wrote the manuscript. All authors read and approved the final version of the manuscript.

## Supplementary information

Additional information is contained in the supplementary information file.

## References

1. Lefevre, T., and F. Thomas. 2008. Behind the scene, something else is pulling the strings: Emphasizing parasitic manipulation in vector-borne diseases. Infection Genetics and Evolution 8: 504–519.

2. Lefèvre, T., J. C. Koella, F. Renaud, H. Hurd, D. G. Biron, and F. Thomas. 2006. New prospects for research on manipulation of insect vectors by pathogens. PLOS Pathog 2: e72.

3. Woolhouse, M. E. J., L. H. Taylor, and D. T. Haydon. 2001. Population biology of multihost pathogens. Science 292: 1109–1112.

4. Heylen, D., H. Sprong, K. van Oers, M. Fonville, H. Leirs, and E. Matthysen. 2014. Are the specialized bird ticks, *Ixodes arboricola* and *I. frontalis*, competent vectors for *Borrelia burgdorferi sensu lato*? Environ Microbiol 16: 1081–1089.

5. Dolan, M. C., J. Piesman, M. L. Mbow, G. O. Maupin, O. Peter, M. Brossard, and W. T. Golde. 1998. Vector competence of *Ixodes scapularis* and *Ixodes ricinus* (Acari: Ixodidae) for three genospecies of *Borrelia burgdorferi*. J Med Entomol 35: 465–470.

6. Zeidner, N. S., L. Gern, J. Piesman, B. S. Schneider, and M. S. Nuncio. 2002. Coinoculation of *Borrelia* spp. with tick salivary gland lysate enhances spirochete load in mice and is tick species-specific. The Journal of parasitology 88: 1276–1278.

7. Ramamoorthi, N., S. Narasimhan, U. Pal, F. K. Bao, X. F. F. Yang, D. Fish, J. Anguita, M. V. Norgard, F. S. Kantor, J. F. Anderson, R. A. Koski, and E. Fikrig. 2005. The Lyme disease agent exploits a tick protein to infect the mammalian host. Nature 436: 573–577.

8. Neelakanta, G., H. Sultana, D. Fish, J. F. Anderson, and E. Fikrig. 2010. *Anaplasma phagocytophilum* induces *Ixodes scapularis* ticks to express an antifreeze glycoprotein gene that enhances their survival in the cold. The Journal of Clinical Investigation 120: 3179–3190.

9. Herrmann, C., and L. Gern. 2015. Search for blood or water is influenced by *Borrelia burgdorferi* in *Ixodes ricinus*. Parasit Vectors 8.

10. Lacroix, R., W. R. Mukabana, L. C. Gouagna, and J. C. Koella. 2005. Malaria infection increases attractiveness of humans to mosquitoes. PLOS Biol 3: e298.

11. Cornet, S., A. Nicot, A. Rivero, and S. Gandon. 2013. Malaria infection increases bird attractiveness to uninfected mosquitoes. Ecol Lett 16: 323–329.

12. De Moraes, C. M., N. M. Stanczyk, H. S. Betz, H. Pulido, D. G. Sim, A. F. Read, and M. C. Mescher. 2014. Malaria-induced changes in host odors enhance mosquito attraction. Proc Natl Acad Sci U S A 111: 11079–11084.

13. Randolph, S. E. 1991. The effect of *Babesia microti* on feeding and survival in its tick vector, *Ixodes trianguliceps*. Parasitology 102: 9–16.

14. Dizij, A., and K. Kurtenbach. 1995. *Clethrionomys glareolus*, but not *Apodemus flavicollis*, acquires resistance to *Ixodes ricinus* L., the main European vector of *Borrelia burgdorferi*. Parasite Immunol (Oxf) 17: 177–183.

15. Ogden, N. H., A. N. J. Casey, N. P. French, J. D. W. Adams, and Z. Woldehiwet. 2002. Field evidence for density-dependent facilitation amongst *Ixodes ricinus* ticks feeding on sheep. Parasitology 124: 117–125.

16. Randolph, S. E. 1979. Population regulation in ticks - role of acquired-resistance in natural and unnatural hosts. Parasitology 79: 141–156.

17. Wikel, S. K., R. N. Ramachandra, D. K. Bergman, T. R. Burkot, and J. Piesman. 1997. Infestation with pathogen-free nymphs of the tick *Ixodes scapularis* induces host resistance to transmission of *Borrelia burgdorferi* by ticks. Infect Immun 65: 335–338.

18. Nazario, S., S. Das, A. M. de Silva, K. Deponte, N. Marcantonio, J. F. Anderson, D. Fish, E. Fikrig, and F. S. Kantor. 1998. Prevention of *Borrelia burgdorferi* transmission in guinea pigs by tick immunity. Am J Trop Med Hyg 58: 780–785.

19. Narasimhan, S., K. DePonte, N. Marcantonio, X. P. Liang, T. E. Royce, K. F. Nelson, C. J. Booth, B. Koski, J. F. Anderson, F. Kantor, and E. Fikrig. 2007. Immunity against *Ixodes scapularis* salivary proteins expressed within 24 hours of attachment thwarts tick feeding and impairs *Borrelia* transmission. PLOS ONE 2.

20. Ribeiro, J. M. C., F. Alarcon-Chaidez, I. M. B. Francischetti, B. J. Mans, T. N. Mather, J. G. Valenzuela, and S. K. Wikel. 2006. An annotated catalog of salivary gland transcripts from *Ixodes scapularis* ticks. Insect Biochem Mol Biol 36: 111–129.

21. Simo, L., M. Kazimirova, J. Richardson, and S. I. Bonnet. 2017. The essential role of tick salivary glands and saliva in tick feeding and pathogen transmission. Front Cell Infect Microbiol 7.

22. Nuttall, P. A., and M. Labuda. 2004. Tick-host interactions: saliva-activated transmission. Parasitology 129: S177–S189.

23. Woldehiwet, Z. 2008. Immune evasion and immunosuppression by *Anaplasma phagocytophilum*, the causative agent of tick-borne fever of ruminants and human granulocytic anaplasmosis. Vet J 175: 37–44.

24. Kurtenbach, K., K. Hanincova, J. I. Tsao, G. Margos, D. Fish, and N. H. Ogden. 2006. Fundamental processes in the evolutionary ecology of Lyme borreliosis. Nat Rev Microbiol 4: 660–669.

25. Stanek, G., and M. Reiter. 2011. The expanding Lyme *Borrelia* complex-clinical significance of genomic species? Clin Microbiol Infec 17: 487–493.

26. Elsner, R. A., C. J. Hastey, K. J. Olsen, and N. Baumgarth. 2015. Suppression of long-lived humoral immunity following *Borrelia burgdorferi* infection. PLOS Pathog 11.

27. Humair, P. F., O. Rais, and L. Gern. 1999. Transmission of *Borrelia afzelii* from *Apodemus* mice and *Clethrionomys* voles to *Ixodes ricinus* ticks: differential transmission pattern and overwintering maintenance. Parasitology 118: 33–42.

28. Talleklint, L., and T. G. T. Jaenson. 1995. Is the small mammal (*Clethrionomys glareolus*) or the tick vector (*Ixodes ricinus*) the primary overwintering reservoir for the Lyme borreliosis spirochete in Sweden. J Wildl Dis 31: 537–540.

29. Kurtenbach, K., A. Dizij, H. M. Seitz, G. Margos, S. E. Moter, M. D. Kramer, R. Wallich, U. E. Schaible, and M. M. Simon. 1994. Differential immune responses to *Borrelia burgdorferi* in European wild rodent species influence spirochete transmission to *Ixodes ricinus* L (Acari, Ixodidae). Infect Immun 62: 5344–5352.

30. van Duijvendijk, G., H. Sprong, and W. Takken. 2015. Multi-trophic interactions driving the transmission cycle of *Borrelia afzelii* between *Ixodes ricinus* and rodents: a review. Parasit Vectors 8: 1–11.

31. Gomez-Chamorro, A., F. Battilotti, C. Cayol, T. Mappes, E. Koskela, N. Boulanger, D. Genné, A. Sarr, and M. J. Voordouw. 2019. Susceptibility to infection with *Borrelia afzelii* and TLR2 polymorphism in a wild reservoir host. Scientific Reports 9: 6711.

32. Belli, A., A. Sarr, O. Rais, R. O. M. Rego, and M. J. Voordouw. 2017. Ticks infected via co-feeding transmission can transmit Lyme borreliosis to vertebrate hosts. Scientific Reports 7: 5006.

33. Jacquet, M., G. Margos, V. Fingerle, and M. J. Voordouw. 2016. Comparison of the lifetime host-to-tick transmission between two strains of the Lyme disease pathogen *Borrelia afzelii*. Parasit Vectors 9.

34. Tonetti, N., M. J. Voordouw, J. Durand, S. Monnier, and L. Gern. 2015. Genetic variation in transmission success of the Lyme borreliosis pathogen *Borrelia afzelii*. Ticks Tick Borne Dis 6: 334–343.

35. Durand, J., M. Jacquet, O. Rais, L. Gern, and M. J. Voordouw. 2017. Fitness estimates from experimental infections predict the long-term strain structure of a vector-borne pathogen in the field. Scientific Reports 7: 1851.

36. Jacquet, M., J. Durand, O. Rais, and M. J. Voordouw. 2015. Cross-reactive acquired immunity influences transmission success of the Lyme disease pathogen, *Borrelia afzelii*. Infection Genetics and Evolution 36: 131–140.

37. Randolph, S. E. 1994. Density-dependent acquired resistance to ticks in natural hosts, independent of concurrent infection with *Babesia microti*. Parasitology 108: 413–419.

38. Gray, G. D., and R. S. Phillips. 1983. Suppression of primary and secondary antibody-responses and inhibition of antigen priming during *Babesia microti* infections in mice. Parasite Immunology 5: 123–134.

39. Vaumourin, E., G. Vourc’h, P. Gasqui, and M. Vayssier-Taussat. 2015. The importance of multiparasitism: examining the consequences of co-infections for human and animal health. Parasit Vectors 8.

40. Porter, J. D., and K. P. McAdam. 1994. The re-emergence of tuberculosis. Annu Rev Public Health 15: 303–323.

41. Walter, K. S., K. M. Pepin, C. T. Webb, H. D. Gaff, P. J. Krause, V. E. Pitzer, and M. A. Diuk-Wasser. 2016. Invasion of two tick-borne diseases across New England: harnessing human surveillance data to capture underlying ecological invasion processes. P Roy Soc B-Biol Sci 283.

42. Diuk-Wasser, M. A., E. Vannier, and P. J. Krause. 2016. Coinfection by *lxodes* tick-borne pathogens: ecological, epidemiological, and clinical consequences. Trends Parasitol 32: 30–42.

43. Andersson, M., K. Scherman, and L. Raberg. 2013. Multiple-strain infections of *Borrelia afzelii*: a role for within-host interactions in the maintenance of antigenic diversity? Am Nat 181: 545–554.

44. Durand, J., C. Herrmann, D. Genné, A. Sarr, L. Gern, and M. J. Voordouw. 2017. Multistrain infections with Lyme borreliosis pathogens in the tick vector. Appl Environ Microbiol 83.

45. Herrmann, C., L. Gern, and M. J. Voordouw. 2013. Species co-occurrence patterns among Lyme borreliosis pathogens in the tick vector *Ixodes ricinus*. Appl Environ Microbiol 79: 7273–7280.

46. Herrmann, C., M. J. Voordouw, and L. Gern. 2013. *Ixodes ricinus* ticks infected with the causative agent of Lyme disease, *Borrelia burgdorferi* sensu lato, have higher energy reserves. Int J Parasitol 43: 477–483.

47. Herrmann, C., and L. Gern. 2012. Do the level of energy reserves, hydration status and *Borrelia* infection influence walking by *Ixodes ricinus* (Acari: Ixodidae) ticks? Parasitology 139: 330–337.

48. Crooks, E., and S. E. Randolph. 2006. Walking by *Ixodes ricinus* ticks: intrinsic and extrinsic factors determine the attraction of moisture or host odour. J Exp Biol 209: 2138–2142.

49. Gray, J. S. 1981. The fecundity of *Ixodes ricinus* (l) (Acarina, Ixodidae) and the mortality of its developmental stages under field conditions. Bull Entomol Res 71: 533–542.

50. Ogden, N. H., L. R. Lindsay, G. Beauchamp, D. Charron, A. Maarouf, C. J. O’Callaghan, D. Waltner-Toews, and I. K. Barker. 2004. Investigating the relationships between temperature and developmental rates of tick *Ixodes scapularis* (Acari: Ixodidae) in the laboratory and field. J Med Entomol 41: 622–633.

51. Ogden, N. H., M. Bigras-Poulin, C. J. O’Callaghan, I. K. Barker, K. Kurtenbach, L. R. Lindsay, and D. Charron. 2007. Vector seasonality, host infection dynamics and fitness of pathogens transmitted by the tick *Ixodes scapularis*. Parasitology 134: 209–227.

52. Dobson, A. D. M., T. J. R. Finnie, and S. E. Randolph. 2011. A modified matrix model to describe the seasonal population ecology of the European tick *Ixodes ricinus*. J Appl Ecol 48: 1017–1028.

53. Gern, L., M. Siegenthaler, C. M. Hu, S. Leuba-Garcia, P. F. Humair, and J. Moret. 1994. *Borrelia burgdorferi* in rodents (*Apodemus flavicollis* and *A. sylvaticus*): Duration and enhancement of infectivity for *Ixodes ricinus* ticks. Eur J Epidemiol 10: 75–80.

54. Richter, D., B. Klug, A. Spielman, and F. R. Matuschka. 2004. Adaptation of diverse Lyme disease spirochetes in a natural rodent reservoir host. Infect Immun 72: 2442–2444.

55. Tsao, J. 2009. Reviewing molecular adaptations of Lyme borreliosis spirochetes in the context of reproductive fitness in natural transmission cycles. Vet Res (Paris) 40.

56. Derdakova, M., V. Dudioak, B. Brei, J. S. Brownstein, I. Schwartz, and D. Fish. 2004. Interaction and transmission of two *Borrelia burgdorferi* sensu stricto strains in a tick-rodent maintenance system. Appl Environ Microbiol 70: 6783–6788.

57. Hanincova, K., N. H. Ogden, M. Diuk-Wasser, C. J. Pappas, R. Iyer, D. Fish, Schwartz, and K. Kurtenbach. 2008. Fitness variation of *Borrelia burgdorferi* sensu stricto strains in mice. Appl Environ Microbiol 74: 153–157.

58. Rynkiewicz, E. C., J. Brown, D. M. Tufts, C.-I. Huang, H. Kampen, S. J. Bent, D. Fish, and M. A. Diuk-Wasser. 2017. Closely-related *Borrelia burgdorferi* (sensu stricto) strains exhibit similar fitness in single infections and asymmetric competition in multiple infections. Parasit Vectors 10: 64.

59. de Roode, J. C., R. Pansini, S. J. Cheesman, M. E. H. Helinski, S. Huijben, A. R. Wargo, A. S. Bell, B. H. K. Chan, D. Walliker, and A. F. Read. 2005. Virulence and competitive ability in genetically diverse malaria infections. Proc Natl Acad Sci U S A 102: 7624–7628.

60. Grillon, A., B. Westermann, P. Cantero, B. Jaulhac, M. J. Voordouw, D. Kapps, E. Collin, C. Barthel, L. Ehret-Sabatier, and N. Boulanger. 2017. Identification of *Borrelia* protein candidates in mouse skin for potential diagnosis of disseminated Lyme borreliosis. Scientific Reports 7: 16719.

61. Kern, A., G. Schnell, Q. Bernard, A. Boeuf, B. Jaulhac, E. Collin, C. Barthel, L. Ehret-Sabatier, and N. Boulanger. 2015. Heterogeneity of *Borrelia burgdorferi* sensu stricto population and its involvement in *Borrelia* pathogenicity: study on murine model with specific emphasis on the skin interface. PLOS ONE 10.

62. Raberg, L. 2012. Infection intensity and infectivity of the tick-borne pathogen *Borrelia afzelii*. J Evol Biol 25: 1448–1453.

63. Wooten, R. M., Y. Ma, R. A. Yoder, J. P. Brown, J. H. Weis, J. F. Zachary, C. J. Kirschning, and J. J. Weis. 2002. Toll-like receptor 2 is required for innate, but not acquired, host defense to *Borrelia burgdorferi*. J Immunol 168: 348–355.

64. Hodzic, E., S. L. Feng, K. J. Freet, and S. W. Barthold. 2003. *Borrelia burgdorferi* population dynamics and prototype gene expression during infection of immunocompetent and immunodeficient mice. Infect Immun 71: 5042–5055.

65. Wang, G., C. Ojaimi, R. Iyer, V. Saksenberg, S. A. McClain, G. P. Wormser, and I. Schwartz. 2001. Impact of genotypic variation of *Borrelia burgdorferi* sensu stricto on kinetics of dissemination and severity of disease in C3H/HeJ mice. Infect Immun 69: 4303–4312.

66. Barthold, S. W. 1999. Specificity of infection-induced immunity among *Borrelia burgdorferi* sensu lato species. Infect Immun 67: 36–42.

67. Connolly, S. E., and J. L. Benach. 2005. The versatile roles of antibodies in *Borrelia* infections. Nat Rev Microbiol 3: 411–420.

68. LaRocca, T. J., and J. L. Benach. 2008. The important and diverse roles of antibodies in the host response to *Borrelia* infections. Curr Top Microbiol Immunol 319: 63–103.

69. Barthold, S. W., E. Hodzic, S. Tunev, and S. Feng. 2006. Antibody-mediated disease remission in the mouse model of Lyme borreliosis. Infect Immun 74: 4817–4825.

70. Liang, F. T., E. L. Brown, T. Wang, R. V. Lozzo, and E. Fikrig. 2004. Protective niche for *Borrelia burgdorferi* to evade humoral immunity. Am J Pathol 165: 977–985.

71. Liang, F., J. Yan, M. L. Mbow, S. Sviat, R. Gilmore, M. Mamula, and E. Fikrig. 2004. *Borrelia burgdorferi* changes its surface antigenic expression in response to host immune responses. Infect Immun 72: 5759–5767.

72. Dunham-Ems, S. M., M. J. Caimano, U. Pal, C. W. Wolgemuth, C. H. Eggers, A. Balic, and J. D. Radolf. 2009. Live imaging reveals a biphasic mode of dissemination of *Borrelia burgdorferi* within ticks. J Clin Invest 119: 3652–3665.

73. Shih, C. M., L. L. Chao, and C. P. Yu. 2002. Chemotactic migration of the Lyme disease spirochete (*Borrelia burgdorferi*) to salivary gland extracts of vector ticks. Am J Trop Med Hyg 66: 616–621.

74. Scheckelhoff, M., S. Telford, M. Wesley, and L. Hu. 2007. *Borrelia burgdorferi* intercepts host hormonal signals to regulate expression of outer surface protein A. Proc Natl Acad Sci U S A 104: 7247–7252.

75. Couret, J., M. C. Dyer, T. N. Mather, S. Han, J. I. Tsao, R. A. Lebrun, and H. S. Ginsberg. 2017. Acquisition of *Borrelia burgdorferi* infection by larval *Ixodes scapularis* (Acari: Ixodidae) associated with engorgement measures. J Med Entomol 54: 1055–1060.

76. Talleklint, L., and T. G. T. Jaenson. 1994. Transmission of *Borrelia burgdorferi* s.l. from mammal reservoirs to the primary vector of Lyme borreliosis, *Ixodes ricinus* (Acari, Ixodidae), in Sweden. J Med Entomol 31: 880–886.

77. Hanincova, K., S. M. Schafer, S. Etti, H. S. Sewell, V. Taragelova, D. Ziak, M. Labuda, and K. Kurtenbach. 2003. Association of *Borrelia afzelii* with rodents in Europe. Parasitology 126: 11–20.

78. Raberg, L., A. Hagstrom, M. Andersson, S. Bartkova, K. Scherman, M. Strandh, and B. Tschirren. 2017. Evolution of antigenic diversity in the tick-transmitted bacterium *Borrelia afzelii*: a role for host specialization? J Evol Biol 30: 1034–1041.

79. Jacquot, M., M. Bisseux, D. Abrial, M. Marsot, E. Ferquel, J. L. Chapuis, G. Vourc’h, and X. Bailly. 2014. High-throughput sequence typing reveals genetic differentiation and host specialization among populations of the *Borrelia burgdorferi* species complex that infect rodents. PLOS ONE 9: e88581.

80. Perez, G., S. Bastian, A. Agoulon, A. Bouju, A. Durand, F. Faille, I. Lebert, Y. Rantier, O. Plantard, and A. Butet. 2016. Effect of landscape features on the relationship between *Ixodes ricinus* ticks and their small mammal hosts. Parasit Vectors 9.

81. Hartemink, N. A., S. E. Randolph, S. A. Davis, and J. A. P. Heesterbeek. 2008. The basic reproduction number for complex disease systems: Defining R-0 for tickborne infections. Am Nat 171: 743–754.

