## Supplemental File for "*Borrelia afzelii* does not suppress the development of anti-tick immunity in bank voles"

#### Section 1 – Phytotron conditions for development of engorged larval ticks

The engorged larval ticks were placed in individual tubes and these tubes were kept in boxes that were stored in a phytotron. The conditions of the phytotron were as follows. Between 5:00–19:00 (14 hours), the light intensity was 3000 lumens and the temperature was 25°C. Between 4:00–5:00 and 19:00–20:00 (2 hours), the light intensity was 1000 lumens and the temperature was 21.5 °C. Between 20:00–04:00 (8 hours), the light intensity was 0 lumen and the temperature was 18°C. The relative humidity was maintained at 85%

#### Section 2 – *B. afzelii* infection status of the bank voles after challenge with uninfected *I. ricinus* nymphs and infected nymphs

Bank voles in the uninfected control group (n = 14) were infested with uninfected *I. ricinus* nymphs whereas bank voles in the infected group (n = 14) were infested with *B. afzelii*-infected *I. ricinus* nymphs. After the nymphal infestation, all bank voles were tested with respect to four infection criteria to determine their actual *B. afzelii* infection status. The four infection criteria were as follows: (1) presence of *B. afzelii*-specific IgG antibodies (optical density of the ELISA > 500 absorbance units), (2) presence of *B. afzelii* in ear tissue biopsy (i.e., spirochete load of *B. afzelii* > 0), (3) presence of *B. afzelii* in xenodiagnostic nymphs (i.e., spirochete load of *B. afzelii* > 0 in at least 1 xenodiagnostic nymph), and (4) presence of live *B. afzelii* spirochetes in culture (i.e. at least 1 culture derived from a xenodiagnostic nymph contains live spirochetes).

The four infection criteria for the 28 bank voles are shown in Table S1. As expected, all of the 14 bank voles in the control group tested negative for the 4 infection criteria (Table S1). Thirteen of the 14 bank voles in the *B. afzelii*-infected group tested positive for 3 or 4 criteria, and these 13 bank voles were therefore considered as infected with *B. afzelii* (Table S1). The bank vole that had been challenged with infected nymphs, but that tested negative for all five criteria was excluded from the analysis. All statistical analyses are therefore based on 13 *B. afzelii*-infected bank voles and 14 uninfected bank voles.

Table S1. The *B. afzelii* infection status is shown for each of the 28 bank voles that belonged to either the *B. afzelii*-infected group or the uninfected control group. The infection status of each bank vole was based on four criteria: (1) *B. afzelii*-specific IgG antibodies, (2) spirochetes in the ear biopsy, (3) *B. afzelii*-infected xenodiagnostic nymphs, and (4) culture of live spirochetes from xenodiagnostic nymphs.

| Vole ID[a] | Treatment[b] | Engorged nymphs[c] | ELISA[d] | Spirochetes in ear biopsy[e] | Xenodiagnosis[f] | Culture of nymphs[g] | Criteria[h] | Infection status[i] |
|---|---|---|---|---|---|---|---|---|
| 193 | Control | 0/3 | 301 | 0 | 0/30 | 0/0 | 0 | Uninfected |
| 195 | Control | 0/4 | 342 | 0 | 0/30 | 0/0 | 0 | Uninfected |
| 196 | Control | 0/4 | 369 | 0 | 0/30 | 0/0 | 0 | Uninfected |
| 197 | Control | 0/1 | 252 | 0 | 0/30 | 0/0 | 0 | Uninfected |
| 198 | Control | 0/3 | 248 | 0 | 0/30 | 0/0 | 0 | Uninfected |
| 199 | Control | 0/4 | 239 | 0 | 0/30 | 0/0 | 0 | Uninfected |
| 206 | Control | 0/4 | 281 | 0 | 0/30 | 0/0 | 0 | Uninfected |
| 209 | Control | 0/4 | 344 | 0 | 0/30 | 0/0 | 0 | Uninfected |
| 211 | Control | 0/3 | 233 | 0 | 0/30 | 0/0 | 0 | Uninfected |
| 216 | Control | 0/2 | 239 | 0 | 0/30 | 0/0 | 0 | Uninfected |
| 224 | Control | 0/5 | 223 | 0 | 0/30 | 0/0 | 0 | Uninfected |
| 228 | Control | 0/2 | 238 | 0 | 0/30 | 0/0 | 0 | Uninfected |
| 230 | Control | 0/5 | 254 | 0 | 0/30 | 0/0 | 0 | Uninfected |
| 231 | Control | 0/3 | 403 | 0 | 0/30 | 0/0 | 0 | Uninfected |
| 192 | Infected | 3/4 (75.0%) | 1120 | 2550 | 4/30 (13.3%) | 2/6 (33.3%) | 4 | Infected |
| 200 | Infected | 1/2 (50.0%) | 5001 | 4315 | 12/27 (44.4%) | 1/6 (16.7%) | 4 | Infected |
| 202 | Infected | 2/2 (100.0%) | 1405 | 21196 | 13/29 (44.8%) | 1/6 (16.7%) | 4 | Infected |
| 204 | Infected | 0/1 (0.0%) | 3865 | 348 | 19/30 (63.3%) | 3/6 (50.0%) | 3 | Infected |
| 205 | Infected | 2/4 (50.0%) | 5475 | 519 | 8/25 (32.0%) | 1/6 (16.7%) | 4 | Infected |
| 214 | Infected | 1/5 (20.0%) | 4309 | 6412 | 2/22 (9.1%) | 0/6 (0.0%) | 3 | Infected |
| 217 | Infected | 4/4 (100.0%) | 3336 | 979 | 7/29 (24.1%) | 2/5 (40.0%) | 4 | Infected |
| 218 | Infected | 1/2 (50.0%) | 2809 | 3704 | 13/27 (44.4%) | 1/4 (25.0%) | 4 | Infected |
| 220 | Infected | 1/3 (33.0%) | 1168 | 17670 | 11/30 (36.7%) | 3/5 (60.0%) | 4 | Infected |
| 222 | Infected | 4/5 (80.0%) | 4096 | 10465 | 11/30 (36.7%) | 1/6 (16.7%) | 4 | Infected |
| 223 | Infected | 0/1 (0.0%) | 218 | 0 | 0/26 (0.0%) | 0/0 (0.0%) | 0 | Uninfected[j] |
| 225 | Infected | 2/3 (66.7%) | 1204 | 6589 | 17/29 (58.6%) | 0/6 (0.0%) | 3 | Infected |
| 226 | Infected | 3/5 (80.0%) | 1587 | 36 | 7/28 (25.0%) | 2/5 (40.0%) | 4 | Infected |
| 229 | Infected | 3/5 (60.0%) | 4011 | 3704 | 12/29 (41.4%) | 1/6 (16.7%) | 4 | Infected |

[a] Vole id is the unique identification number assigned to each bank vole in the study

[b] Treatment refers to whether the bank vole was randomly assigned to the *B. afzelii*-infected group or the uninfected control group.

[c] Engorged nymphs is the number of engorged challenge nymphs that tested positive for *B. afzelii* compared to the total number of engorged challenge nymphs that were collected. The percentage of engorged challenge nymphs that tested positive for *B. afzelii* is shown in brackets.

[d] ELISA is the strength of the *Borrelia afzelii*-specific IgG antibody response as measured by the commercial ELISA assay. The optical density (OD) was measured every 2 minutes over a period of 60 minutes. The area under the curve of OD versus time was integrated to give the values shown. Individuals with an optical density > 500 units are considered infected.

[e] Spirochetes in ear biopsy is the number of spirochetes in the ear tissue biopsy (2 mm diameter) as estimated by our qPCR assay. Individuals with a spirochete load > 1 per ear tissue biopsy are considered infected.

[f] Xenodiagnosis refers to the xenodiagnostic larvae that fed on the bank voles and that subsequently molted into xenodiagnostic nymphs. The number of xenodiagnostic nymphs that tested positive for *B. afzelii* compared to the total number of xenodiagnostic nymphs that were tested using qPCR are shown. The percentage of xenodiagnostic nymphs that tested positive for *B. afzelii* is shown in brackets. Individuals that infect > 0% of the xenodiagnostic nymphs are considered infected.

[g] Culture is the number of xenodiagnostic nymphs that yielded a live spirochete culture compared to the total number of xenodiagnostic nymphs that were placed into BSK media. Individuals that produce at least 1 live spirochete culture are considered infected.

[h] Criteria is the number of infection status criteria that were met by each bank vole and ranges from 0 to 4.

[i] Infection status is whether a vole was considered to be infected with *B. afzelii* or not.

[j] Bank vole 223 did not become infected following exposure to *B. afzelii*-infected nymphs.

#### Section 3 – Relationship between engorged larval weight and flat nymphal weight

For a sample of 880 *I. ricinus* ticks, the engorged larval weight (μg) and the unfed nymphal weight (μg) were measured for each individual tick. A Pearson correlation test was used to test whether there was a relationship between the engorged larval weight and the flat nymphal weight. The engorged larval weight was strongly and positively correlated with the flat nymphal weight (Figure 3; Pearson correlation: $r = 0.796$, $df = 879$, $t = 38.943$, $p < 0.001$). This result shows that the weight of the engorged larva influences the weight of the resultant flat nymph.

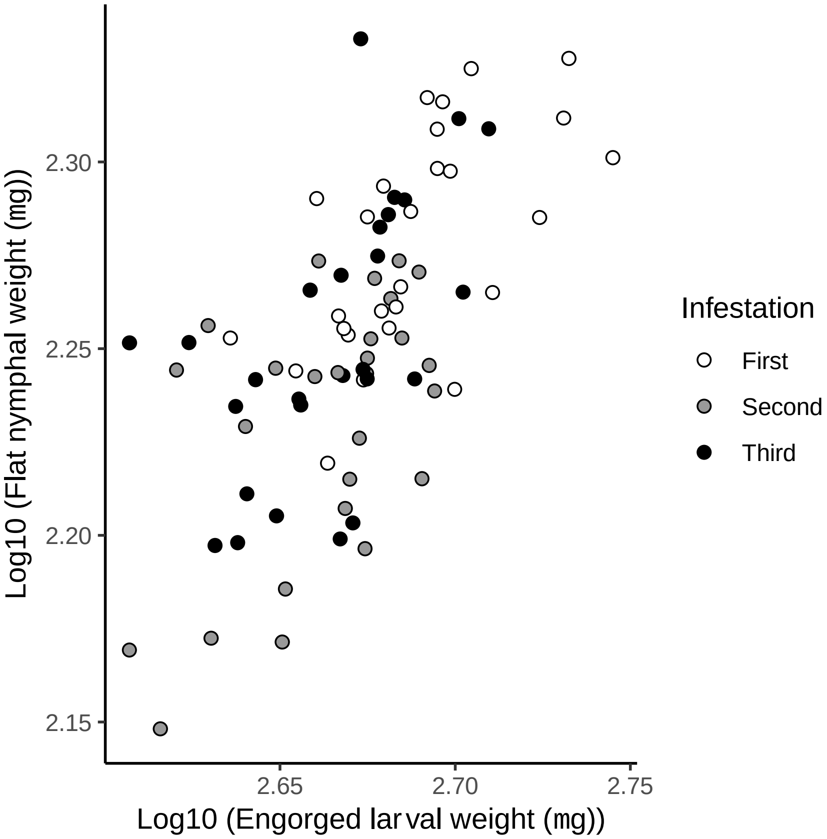

Figure S1. There is a strong and positive relationship between the weight of the engorged larvae (μg) and the weight of the resultant flat nymphs (μg). Larval ticks that take a larger blood meal molt into larger nymphs. The correlation between engorged larval weight and flat nymphal weight was highly significant ($r = 0.796$, $df = 879$, $t = 38.943$, $p < 0.001$).

#### Section 4 – Additional analysis of the tick phenotypes

In the main manuscript, we tested whether the bank voles developed acquired immunity against *I. ricinus* ticks over the three successive infestations and whether infection with *B. afzelii* influenced the development of this anti-tick immunity. The four tick phenotypes that were analyzed included engorged larval weight, flat nymphal weight, larva-to-nymph molting time, and larva-to-nymph molting success. Each of these four tick phenotypes was analyzed as a function of two fixed factors: *B. afzelii* infection status, infestation number, and their interaction using linear mixed effects models (LMMs) or generalized linear mixed effects models (GLMMs). Here we present the parameter estimates of the LMMs and the GLMMs for each of the four tick phenotypes. In Tables S2, S3, S4, S5, S6, S7, S8, and S9 we show the parameter estimates from the most parsimonious model for each tick phenotype.

Table S2A. LMM parameter estimates are shown for the main effects and interaction of infection status and infestation on the log10-transformed weight of the engorged *I. ricinus* larvae. The engorged larval weight was measured in µg. The intercept refers to the log10-transformed engorged weight of larval ticks that fed on uninfected bank voles during the first infestation.

| Fixed effects | Estimate | Standard error | t-value | p-value |
|---|---|---|---|---|
| (Intercept) | 2.682 | 0.005 | 489.314 | < 0.001 |
| 2nd – 1st infestation | -0.019 | 0.007 | -2.747 | 0.006 |
| 3rd – 1st infestation | -0.029 | 0.007 | -4.373 | < 0.001 |
| Infected – Control | 0.005 | 0.008 | 0.620 | 0.537 |
| 2nd – 1st infestation\| Infected | -0.021 | 0.010 | -2.175 | 0.030 |
| 3rd – 1st infestation\| Infected | 0.006 | 0.009 | 0.691 | 0.490 |

Table S2B. LMM parameter estimates are shown for the main effects of infection status and infestation on the log10-transformed weight of the engorged *I. ricinus* larvae. The engorged larval weight was measured in µg. The intercept refers to the log10-transformed engorged weight of larval ticks that fed on uninfected bank voles during the first infestation.

| Fixed effects | Estimate | Standard error | t-value | p-value |
|---|---|---|---|---|
| (Intercept) | 2.684 | 0.005 | 549.492 | < 0.001 |
| 2nd – 1st infestation | -0.029 | 0.005 | -6.044 | < 0.001 |
| 3rd – 1st infestation | -0.025 | 0.005 | -5.464 | < 0.001 |
| Infected – Control | 0.001 | 0.006 | 0.169 | 0.867 |

Table S3. The parameter estimates of the weight of the engorged *I. ricinus* larvae are shown for each of the six combinations of infection status and infestation. Engorged larval weight is measured in µg. The means are shown with their 95% confidence intervals where LL and UL refer to the lower limit and upper limit, respectively.

| Infection Status | Infestation | Larval weight | 95% LL | 95% UL |
|---|---|---|---|---|
| Control | First | 480.84 | 469.32 | 492.86 |
| Control | Second | 460.26 | 448.37 | 473.39 |
| Control | Third | 449.78 | 438.77 | 462.05 |
| Infected | First | 486.41 | 474.39 | 498.59 |
| Infected | Second | 443.61 | 432.15 | 456.08 |
| Infected | Third | 462.38 | 450.58 | 473.77 |

Table S4. LMM parameter estimates are shown for the main effects of infection status and infestation on the log10-transformed weight of the unfed *I. ricinus* nymphs. The unfed nymphal weight was measured in µg. The intercept refers to the log10-transformed weight of unfed nymphs that fed as larvae on uninfected bank voles during the first infestation.

| Fixed effects | Estimate | Standard error | t-value | p-value |
|---|---|---|---|---|
| (Intercept) | 2.270 | 0.007 | 348.258 | < 0.001 |
| $2^{nd}$ – $1^{st}$ infestation | -0.052 | 0.006 | -8.731 | < 0.001 |
| $3^{rd}$ – $1^{st}$ infestation | -0.026 | 0.006 | -4.361 | < 0.001 |
| Infected – Control | 0.006 | 0.008 | 0.782 | 0.441 |

Table S5. The parameter estimates of the weight of the unfed *I. ricinus* nymphs are shown for of the six combinations of infection status and infestation. Unfed nymphal weight is measured in µg. The means are shown with their 95% confidence intervals where LL and UL refer to the lower limit and upper limit, respectively.

| Infection Status | Infestation | Nymphal weight | 95% LL | 95% UL |
|---|---|---|---|---|
| Control | First | 187.37 | 181.50 | 193.43 |
| Control | Second | 166.82 | 160.88 | 172.99 |
| Control | Third | 171.82 | 165.75 | 178.12 |
| Infected | First | 187.77 | 181.87 | 193.87 |
| Infected | Second | 165.61 | 159.70 | 171.74 |
| Infected | Third | 181.64 | 175.24 | 188.28 |

Table S6. LMM parameter estimates are shown for the main effects of infection status and infestation on the larva-to-nymph molting time of *I. ricinus*. The larva-to-nymph molting time is measured in days. The intercept refers to the larva-to-nymph molting time of larvae that fed on uninfected bank voles during the first infestation.

| Fixed effects | Estimate | Standard error | t-value | p-value |
|---|---|---|---|---|
| (Intercept) | 51.261 | 1.134 | 45.191 | < 0.001 |
| 2nd – 1st infestation | -4.046 | 1.134 | -3.568 | < 0.001 |
| 3rd – 1st infestation | -16.256 | 1.018 | -15.973 | < 0.001 |
| Infected – Control | -0.080 | 1.489 | -0.054 | 0.958 |

Table S7. The parameter estimates of the molting time are shown for each infestation. The infestation had a significant effect on the larva-to-nymph molting time. The molting time refers to the number of days it takes for an engorged larval tick to molt into a nymphal tick. The means are shown with their 95% confidence intervals where LL and UL refer to the lower limit and upper limit, respectively.

| Infestation | Molting time | 95% LL | 95% UL |
|---|---|---|---|
| First | 51.22 | 49.55 | 52.89 |
| Second | 47.18 | 44.99 | 49.37 |
| Third | 34.96 | 33.00 | 36.93 |

Table S8. GLMM parameter estimates are shown for the main effects of infection status and infestation on the larva-to-nymph molting success of *I. ricinus*. Larva-to-nymph molting success is a binomial variable (Not molted = 0, Molted = 1). The intercept refers to the molting success of the larvae that fed on uninfected bank voles during the first infestation.

| Fixed effects | Estimate | Standard error | z-value | p-value |
|---|---|---|---|---|
| (Intercept) | 2.055 | 0.112 | 18.269 | < 0.001 |
| 2nd – 1st infestation | -0.023 | 0.156 | -0.148 | 0.883 |
| 3rd – 1st infestation | -0.690 | 0.123 | -5.633 | < 0.001 |
| Infected – Control | -0.158 | 0.128 | -1.241 | 0.215 |

Table S9. The parameter estimates of the molting success are shown for each infestation. The infestation had a significant effect on the molting success. The larva-to-nymph molting success refers to the percentage of engorged larval ticks that developed into the nymphal stage. The means are shown with their 95% confidence intervals where LL and UL refer to the lower limit and upper limit, respectively.

| Infestation | Molting success (%) | 95% LL | 95% UL |
|---|---|---|---|
| First | 87.7 | 85.6 | 89.9 |
| Second | 87.4 | 84.2 | 90.5 |
| Third | 78.1 | 75.4 | 80.8 |

#### Section 5 – The spirochete load of *B. afzelii* in the ear tissues of the bank voles decreased over the three infestations

The spirochete loads in the bank vole ear biopsies (2 mm diameter) decreased over the three infestations (Figure S2) and this effect was statistically significant (LLR test: $\chi_2$ = 38.231, df = 2, p < 0.001). For the first, second, and third larval infestation, the mean spirochete load in the ear biopsy (measured in number of spirochetes) was 5883 (95% CI: 1516–22834), 202 (95% CI: 52–786), and 13 (95% CI: 3–50), respectively.

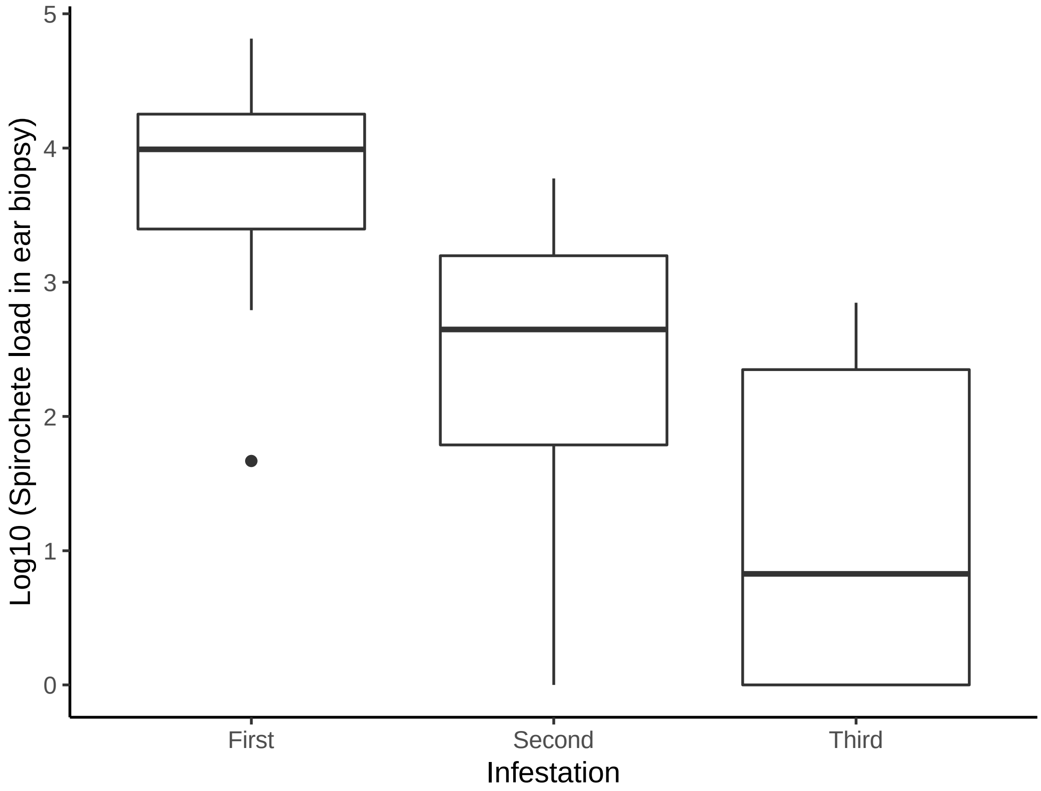

Figure S2. The spirochete load of *B. afzelii* in the ear biopsies of the bank voles decreased over the three successive infestations. The spirochete load is the number of *B. afzelii* spirochetes in the whole ear tissue biopsy (2 mm diameter). Each of the infected bank voles (n = 13) was biopsied at 26, 51, and 106 days PI. Shown are the medians (black line), the 25th and 75th percentiles (edges of the box), the minimum and maximum values (whiskers), and the outliers (circles).

#### Section 6 – Host-to-tick transmission of *B. afzelii* from infected bank voles to *I. ricinus* ticks decreased over the three infestations

Host-to-tick transmission decreased over the three infestations (Figure S3) and this effect was statistically significant (LLR test: $\chi_2 = 41.808$, df = 2, $p < 0.001$). For the first, second, and third larval infestations, the host-to-tick transmission was 59.4% (76/128), 35.6% (42/118), and 20.3% (26/128), respectively.

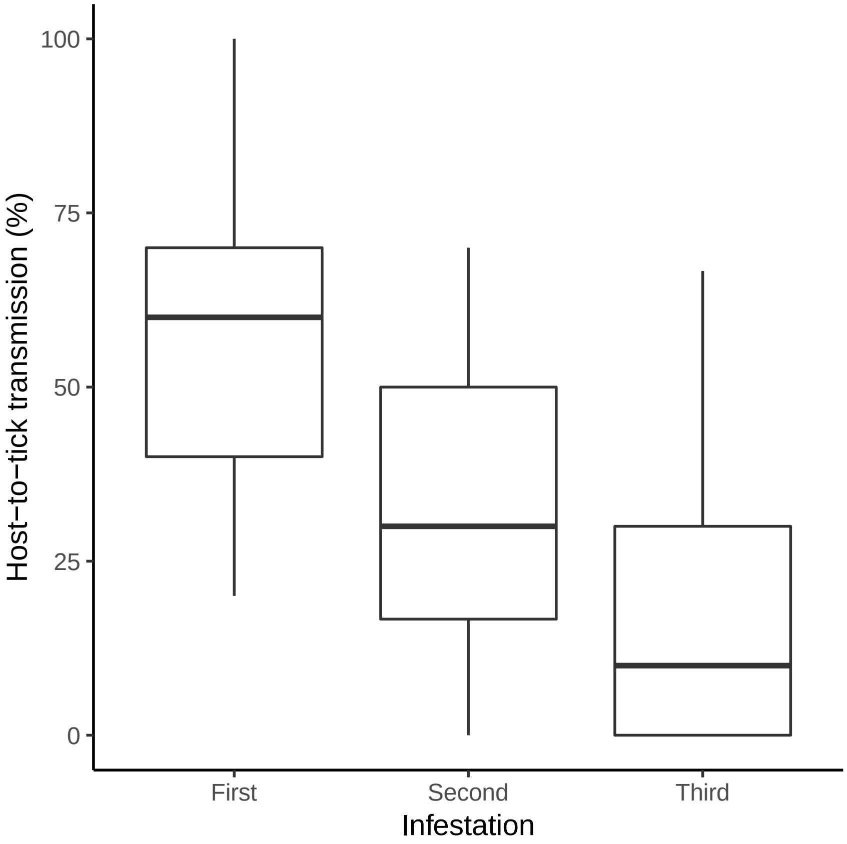

Figure S3. The host-to-tick transmission of *B. afzelii* from the infected bank voles to the *I. ricinus* ticks decreased over the three successive infestations. Host-to-tick transmission refers to the percentage of nymphs that acquired the *B. afzelii* infection during the larval blood meal. Each of the infected bank voles (n = 13) was infested with 50–100 larval *I. ricinus* ticks at 27, 55, and 84 days PI. The engorged larval ticks were allowed to molt into nymphs and the percentage of *B. afzelii*-infected nymphs was estimated using qPCR. Each host-to-tick transmission data point is based on 10 nymphs. Shown are the medians (black line), the 25th and 75th percentiles (edges of the box), the minimum and maximum values (whiskers), and the outliers (circles).

#### Section 7 – Analysis of the spirochete load of *B. afzelii* in the *I. ricinus* nymphs that had fed as larval ticks on the infected bank voles

We modeled the spirochete load in the flat nymphs as a function of the tick SGE-specific IgG antibody response, the spirochete load in the ear biopsy, and host-to-tick transmission. The mean spirochete load in the nymphs decreased over the three infestations and this effect was statistically significant (LMM: $\chi_2$ = 7.807, df = 2, p = 0.020). For the first, second, and third larval infestation, the mean spirochete load in the nymph was 608.9 (95% CI: 241.8–824.4), 234.1 (95% CI: 84.8–302.9), and 455.1 (95% CI: 174.8–743.9), respectively.

The tick SGE-specific IgG antibody response (LLR: $\chi_2$ = 0.089, df = 1, p = 0.766) and the ear spirochete load (LLR: $\chi_2$ = 0.073, df = 1, p = 0.787) had no effect on the spirochete load in the nymphs. Host-to-tick transmission had a significant positive effect on the mean spirochete load in the nymphs (Figure S4; LLR: $\chi_2$ = 6.246, df = 1, p = 0.012; slope = 1.12 ± 0.450).

The positive relationship between host-to-tick transmission and the nymphal spirochete load occurred because both of these two variables decreased over the three infestations. One explanation is that acquired anti-tick immunity created an increasingly hostile environment for the *B. afzelii* spirochetes in the feeding lesion of the larval ticks. This anti-tick immunity would have reduced the probability of host-to-tick transmission and spirochete growth in the subset of larval ticks that did become infected with *B. afzelii*. These results are consistent with previous studies that found a positive relationship between host-to-tick transmission and the nymphal spirochete load of *B. afzelii* (1-3). These results are also consistent with two recent studies that found that the spirochete load of *B. afzelii* was much lower in nymphs that had fed as larvae under conditions that induced a stronger anti-tick response in the rodent host (4, 5). The nymphal spirochete load may have important consequences for tick-to-host transmission (6, 7). We recently showed that *ospC* strains of *B. afzelii* that establish a high spirochete load in field-collected *I. ricinus* nymphs have a higher frequency in these populations of *I. ricinus* (7) and we suggested that such strains have higher nymph-to-host transmission of *B. afzelii*. In contrast, a recent lab study showed that a 20-fold difference in nymphal spirochete load had no effect on the probability of nymph-to-host transmission of *B. afzelii* (4). Thus, the importance of nymphal spirochete load for nymph-to-host transmission of *B. burgdorferi* sl pathogens is currently not clear.

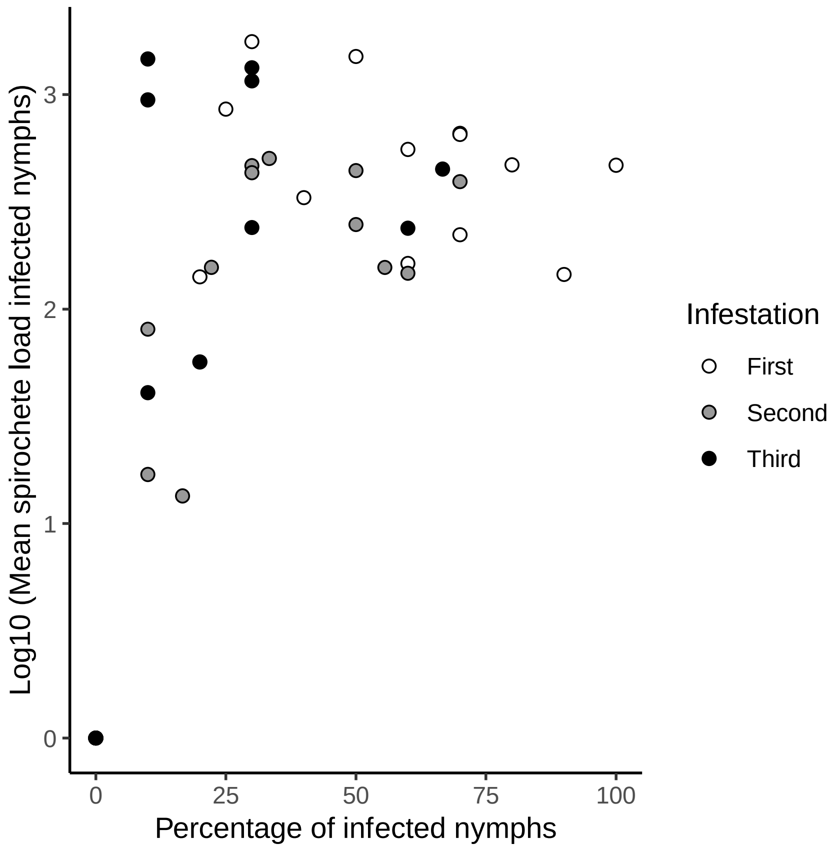

Figure S4. The mean spirochete load in the *B. afzelii*-infected nymphs is positively correlated with host-to-tick transmission. Data are shown for the subset of *B. afzelii*-infected bank voles (n = 13 individuals) at the time of the first (open white circles), second (solid grey circles), and third infestation (solid black circles). The bank voles were infested with larval ticks at 27, 55, and 84 days PI. The 39 data points represent the 13 *B. afzelii*-infected bank voles at each of the three infestations. Spirochete loads are calculated for the entire nymph.

### Section 8 – Analysis of the spirochete load of *B. afzelii* in the organ tissue samples of the bank voles at 106 days post-infection

**Materials and methods**

The dramatic decline in the spirochete load of *B. afzelii* in the ear tissue biopsies (Figure S1) led us to suspect that the bank voles were clearing their infections. We therefore decided to test the *B. afzelii* infection status of various organs after the bank voles were euthanized. There were 28 bank voles that survived to the end of the experiment: 14 in the *B. afzelii*-infected group (of which 13 were truly infected) and 14 in the uninfected control group. All 28 bank voles were sacrificed at 106 days post-infection (PI) via asphyxiation with $CO_2$. The bank voles were dissected under sterile conditions and the following five organs were removed: ear, bladder, heart, ventral skin, and dorsal skin. The ear was sampled using an ear punch (2 mm diameter). For the bladder, heart, ventral skin, and dorsal skin, ~20 to 25 mg of tissue were obtained. The dissection equipment was sterilized with bleach in between individuals and in between organs of the same individual to avoid contamination. A total of 140 bank vole tissue samples were obtained (28 bank voles* 5 tissue samples per bank vole = 140 tissue samples).

The DNA was extracted from 140 tissue samples using the QIAGEN DNeasy® Blood and Tissue Kit and following the manufacturer's protocol. For the ear, the whole biopsy (2 mm diameter) was used for DNA extraction. For the bladder, heart, ventral skin, and dorsal skin, we obtained the wet weight of each tissue sample using a balance before DNA extraction. The DNA concentration of each DNA extraction was measured using a Nanodrop. The spirochete load of the tissue samples was estimated using a qPCR that amplifies a 132 bp fragment of the *flagellin* gene of *B. afzelii*. Each qPCR contained 84 samples from individual bank voles, triplicates of the standards (10, 100, 1000, 10000 flagellin gene copies) and negative controls ($H_20$). For each tissue sample, we standardized the spirochete load in two different ways. The number of spirochetes in the DNA extraction was divided by the amount of tissue that was extracted to calculate the number of spirochetes per mg of tissue. The numbers of spirochetes in 1 µl of DNA extract were divided by the DNA concentration of the extract to calculate the number of spirochetes per mg of DNA. The spirochete density in rodent tissues is low so that the DNA concentration of *B. burgdorferi* sl-infected rodent tissue is mostly a measure of rodent DNA (and hence rodent cell content).

**Statistical analysis**

We standardized the spirochete load in the bank vole tissue samples in two different ways: relative to the mass of the tissue sample and relative to the DNA concentration of the tissue sample. For the subset of *B. afzelii*-infected bank voles (n = 13), we used a Pearson correlation test to determine whether there was a significant correlation between these two different ways to standardize the spirochete load.

To test whether there were differences in the probability of infection between the five organs, we used a GLMM with binomial errors for the subset of *B. afzelii*-infected bank voles (n = 13). In this model, the binomial response variable was whether the organ tissue sample was uninfected (0) or infected (1). Organ was modeled as a fixed factor and the bank vole ID was used as a random factor.

To test whether there were differences in tissue spirochete load between the five organs, we used an LMM with normal errors for the subset of *B. afzelii*-infected bank voles (n = 13). In this model, the response variable was the standardized spirochete load, which was transformed using log10(spirochete load + 1), to normalize the residuals. Organ was modeled as a fixed factor and bank vole ID as a random factor.

In addition, we wanted to test whether the spirochete loads in some organs, such as the skin, are more important for host-to-tick transmission than other organs. We had estimated the spirochete load at 106 days PI in five organs: bladder, ear, heart, dorsal skin, and ventral skin. For each organ, the spirochete loads were expressed as the number of spirochetes per mg of tissue or as the number of spirochetes per mg of DNA. The spirochete loads were log10(X + 1)-transformed. For each infected bank vole, the lifetime host-to-tick transmission was calculated as the percentage of *B. afzelii*-infected nymphs summed over the three infestations. We modelled the lifetime host-to-tick transmission as a function of the spirochete loads in five organs: heart, bladder, ear, dorsal skin, and ventral skin using a generalized linear model (GLM) with binomial errors.

**Results**

The analysis of the tissue spirochete loads of the organs further confirmed the infection status of the 28 bank voles. In the control group, 0 of 14 bank voles and 0 of the 70 tissue samples tested positive for *B. afzelii* (Tables S10 and S11). In the *B. afzelii*-infected group, 12 the 14 bank voles and 39 of the 70 tissue samples tested positive for *B. afzelii* (Tables S10 and S11). The two bank voles in the *B. afzelii*-infected group that tested negative for *B. afzelii* for all 5 tissues samples were bank voles 223 and 205 (Tables S10 and S11). According to our 4 other infection criteria (Table S1), bank voles 223 and 205 were uninfected and infected, respectively.

Figure S3 shows that the two methods of standardizing the tissue spirochete load give very similar results. The correlation between the two methods of standardizing the spirochete load was highly significant (Figure S3; $r = 0.989$, $t = 54.043$, $df = 63$, $p < 2.2e-16$).

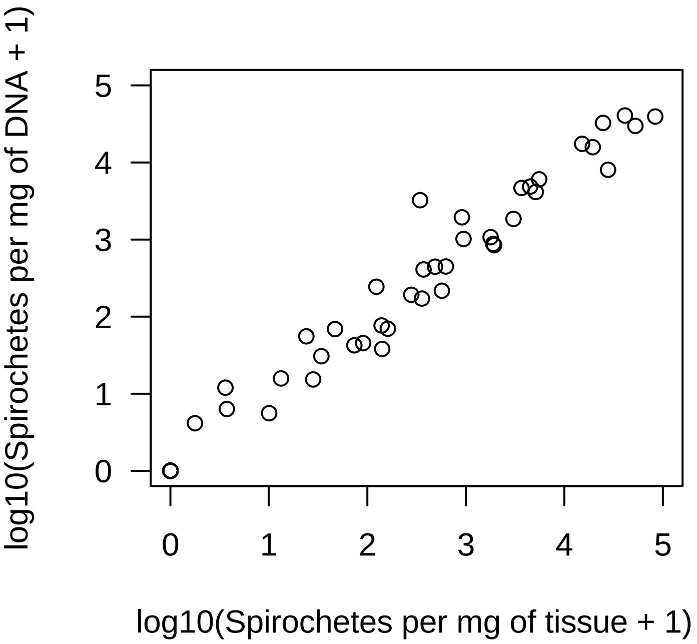

Figure S5. The two methods of standardizing the spirochete load in the bank vole tissues give very similar results. The tissue spirochete loads were expressed as the spirochetes per mg of tissue (horizontal axis) or the spirochetes per mg of DNA (vertical axis).

Organ had a highly significant effect on the probability of whether the tissue sample tested positive for *B. afzelii* in the qPCR ($\Delta \chi_2$ = 34.93, $\Delta$ df =4, p = 4.802e-07). The percentage of infected tissue samples was ranked as follows: 15.4% (2/13) in the heart, 38.4% (5/13) in the ear, 61.5% (8/13) in the bladder, 92.3% (12/13) in the belly skin, and 92.3% (12/13) in the neck skin.

Organ had a highly significant effect on the spirochete load per mg of tissue ($\Delta \chi_2$ = 53.271, $\Delta$ df =4, p = 7.479e-11) and on the spirochete load per mg of DNA. ($\Delta \chi_2$ = 46.724, $\Delta$ df =4, p = 1.741e-09). The mean spirochete load was ranked from lowest to highest as follows: heart, ear, bladder, neck skin, and belly skin (Table S7). The mean spirochete load in the belly skin was ~1000 times higher compared to the heart. Interestingly, the spirochete load was highest in the skin (Table S7), which is the organ that is most important for host-to-tick transmission.

Table S6. Mean tissue spirochete loads of *B. afzelii* differ among the five organs in the subset of infected bank voles. The spirochete loads were standardized per mg of tissue and per mg of DNA. Shown are the means and the lower limit (LL) and upper limit (UL) of the 95% confidence interval.

| Organ | Standardization | Mean | 95% LL | 95% UL |
|---|---|---|---|---|
| Bladder | Mg of Tissue | 16.7 | 3.8 | 74.6 |
| Ear | Mg of Tissue | 10.2 | 2.3 | 45.4 |
| Heart | Mg of Tissue | 1.2 | 0.3 | 5.2 |
| Skin belly | Mg of Tissue | 1413.5 | 317.0 | 6302.0 |
| Skin neck | Mg of Tissue | 490.9 | 110.1 | 2188.5 |
| Bladder | Mg of DNA | 22.3 | 4.9 | 102.2 |
| Ear | Mg of DNA | 10.3 | 2.2 | 47.0 |
| Heart | Mg of DNA | 1.3 | 0.3 | 5.9 |
| Skin belly | Mg of DNA | 1151.3 | 251.4 | 5271.9 |
| Skin neck | Mg of DNA | 278.4 | 60.8 | 1274.7 |

The correlations in spirochete load were all positive, except the correlation between heart and ear, but none were statistically significant, except the correlation between the belly skin and the neck skin ($r = 0.728$, $df = 11$, $p = 0.005$; Table S8). The GLM analysis of lifetime host-to-tick transmission as a function of the spirochete loads in the five different organs (heart, bladder, ear, belly skin, and neck skin) was overdispersed (residual deviance = 22.630, residual degrees of freedom = 7), and the data was therefore re-analyzed using a quasibinomial distribution. For the spirochete loads that were standardized to the DNA concentration, the spirochete load in the belly skin was positively related to the lifetime host-to-tick transmission, but the slope was not significantly different from zero (Table S9; $p = 0.0738$). For the spirochete loads that were standardized to the weight of the tissue sample, the spirochete load in the belly skin was positively related to the lifetime host-to-tick transmission, but the slope was not significantly different from zero (Table S9; $p = 0.0551$).

Table S7. Correlation matrix of the tissue spirochete loads between the five organs. The five organs include: bladder, ear, heart, belly skin, and neck skin. Data are based on the subset of 13 bank voles that were infected with *B. afzelii*. All tissue spirochete loads were log10(X + 1)-transformed to improve their fit to the normal distribution. The Pearson correlation coefficients are shown on the right side of the diagonal and the p-values of the correlation on the left side of the diagonal.

| Organ | Bladder | Ear | Heart | Belly skin | Neck skin |
|---|---|---|---|---|---|
| Bladder | *** | 0.312 | 0.360 | 0.508 | 0.393 |
| Ear | 0.299 | *** | -0.053 | 0.428 | 0.164 |
| Heart | 0.227 | 0.862 | *** | 0.317 | 0.215 |
| Belly skin | 0.077 | 0.144 | 0.291 | *** | **0.728** |
| Neck skin | 0.185 | 0.592 | 0.481 | **0.005** | *** |

Table S8. The mean spirochete loads in five different organs do not influence the lifetime host-to-tick transmission. The five organs include: bladder, ear, heart, belly skin, and neck skin. The mean tissue spirochete load for each organ was standardized by either the DNA concentration of the tissue DNA extraction (units are spirochetes per mg of DNA) or the weight of the tissue sample (units are spirochetes per mg of tissue).

| Correction | Organ | Slope | S.E. | t-statistic | P |
|---|---|---|---|---|---|
| DNA | Intercept | -0.91479 | 0.51333 | -1.782 | 0.1179 |
| DNA | Log10(Bladder+1) | -0.07309 | 0.17752 | -0.412 | 0.6929 |
| DNA | Log10(Ear+1) | -0.20156 | 0.16539 | -1.219 | 0.2624 |
| DNA | Log10(Heart+1) | -0.14006 | 0.81856 | -0.171 | 0.869 |
| DNA | Log10(Belly skin+1) | 0.47081 | 0.22409 | 2.101 | 0.0738 |
| DNA | Log10(Neck skin+1) | -0.31438 | 0.26042 | -1.207 | 0.2666 |
| Tissue | Intercept | -0.9774 | 0.58363 | -1.675 | 0.1379 |
| Tissue | Log10(Bladder+1) | -0.07508 | 0.19189 | -0.391 | 0.7072 |
| Tissue | Log10(Ear+1) | -0.26381 | 0.17887 | -1.475 | 0.1837 |
| Tissue | Log10(Heart+1) | -0.43307 | 1.25817 | -0.344 | 0.7408 |
| Tissue | Log10(Belly skin+1) | 0.56235 | 0.28186 | 1.995 | 0.0862 |
| Tissue | Log10(Neck skin+1) | -0.34201 | 0.29665 | -1.153 | 0.2868 |

Table S8. The *B. afzelii* spirochete loads are shown for the dissected organs of the bank voles. The units of the spirochete load are the number of spirochetes per mg of DNA.

| Vole ID[a] | Sex[b] | Treatment[c] | N positive[d] | Bladder[e] | Ear[f] | Heart[g] | Skin belly[h] | Skin neck[i] | Status[j] |
|---|---|---|---|---|---|---|---|---|---|
| 193 | F | Control | 0 | 0 | 0 | 0 | 0 | 0 | Uninfected |
| 195 | F | Control | 0 | 0 | 0 | 0 | 0 | 0 | Uninfected |
| 196 | F | Control | 0 | 0 | 0 | 0 | 0 | 0 | Uninfected |
| 197 | F | Control | 0 | 0 | 0 | 0 | 0 | 0 | Uninfected |
| 198 | M | Control | 0 | 0 | 0 | 0 | 0 | 0 | Uninfected |
| 199 | M | Control | 0 | 0 | 0 | 0 | 0 | 0 | Uninfected |
| 206 | F | Control | 0 | 0 | 0 | 0 | 0 | 0 | Uninfected |
| 209 | F | Control | 0 | 0 | 0 | 0 | 0 | 0 | Uninfected |
| 211 | F | Control | 0 | 0 | 0 | 0 | 0 | 0 | Uninfected |
| 216 | M | Control | 0 | 0 | 0 | 0 | 0 | 0 | Uninfected |
| 224 | F | Control | 0 | 0 | 0 | 0 | 0 | 0 | Uninfected |
| 228 | F | Control | 0 | 0 | 0 | 0 | 0 | 0 | Uninfected |
| 230 | M | Control | 0 | 0 | 0 | 0 | 0 | 0 | Uninfected |
| 231 | M | Control | 0 | 0 | 0 | 0 | 0 | 0 | Uninfected |
| 192 | F | Infected | 3 | 41 | 0 | 0 | 37 | 872 | Infected |
| 200 | F | Infected | 3 | 1848 | 0 | 0 | 4647 | 1070 | Infected |
| 202 | M | Infected | 3 | 68 | 0 | 0 | 17450 | 170 | Infected |
| 204 | M | Infected | 4 | 11 | 0 | 3 | 6039 | 68 | Infected |
| 205 | M | Infected | 0 | 0 | 0 | 0 | 0 | 0 | Infected |
| 214 | F | Infected | 2 | 0 | 0 | 0 | 44 | 75 | Infected |
| 217 | NA | Infected | 3 | 0 | 446 | 0 | 4845 | 190 | Infected |
| 218 | F | Infected | 4 | 1939 | 4126 | 0 | 29764 | 216 | Infected |
| 220 | M | Infected | 4 | 54 | 243 | 0 | 442 | 408 | Infected |
| 222 | F | Infected | 4 | 14 | 1015 | 0 | 15767 | 844 | Infected |
| 223 | M | Infected | 0 | 0 | 0 | 0 | 0 | 0 | Uninfected[k] |
| 225 | M | Infected | 2 | 0 | 0 | 0 | 40599 | 39389 | Infected |
| 226 | M | Infected | 5 | 3230 | 30 | 5 | 32525 | 8011 | Infected |
| 229 | M | Infected | 2 | 0 | 0 | 0 | 5 | 15 | Infected |

[a] Vole ID is the unique identification number assigned to each bank vole in the study.
[b] Sex refers to whether the bank vole is female (F) or male (M).
[c] Treatment refers to whether the bank vole was randomly assigned to the *B. afzelii*-infected group or the uninfected control group.
[d] N positive is the number of organs that tested positive for *B. afzelii* for each bank vole.
[e] Bladder is the number of spirochetes per mg of DNA in the bladder as estimated by our qPCR assay.
[f] Ear is the number of spirochetes per mg of DNA in the ear as estimated by our qPCR assay.
[g] Heart is the number of spirochetes per mg of DNA in the heart as estimated by our qPCR assay.
[h] Skin belly is the number of spirochetes per mg of DNA in the belly skin as estimated by our qPCR assay.
[i] Skin neck is the number of spirochetes per mg of DNA in the neck skin as estimated by our qPCR assay.
[j] Infection status is whether a vole was considered to be infected with *B. afzelii* or not (from Table S1).
[k] Bank vole 223 did not become infected following exposure to *B. afzelii*-infected ticks.

Table S9. The *B. afzelii* spirochete loads are shown for the dissected organs of the bank voles. The units of the spirochete load are the number of spirochetes per mg of tissue.

| Vole ID[a] | Sex[b] | Treatment[c] | N positive[d] | Bladder[e] | Ear[f] | Heart[g] | Skin belly[h] | Skin neck[i] | Status[j] |
|---|---|---|---|---|---|---|---|---|---|
| 193 | F | Control | 0 | 0 | 0 | 0 | 0 | 0 | Uninfected |
| 195 | F | Control | 0 | 0 | 0 | 0 | 0 | 0 | Uninfected |
| 196 | F | Control | 0 | 0 | 0 | 0 | 0 | 0 | Uninfected |
| 197 | F | Control | 0 | 0 | 0 | 0 | 0 | 0 | Uninfected |
| 198 | M | Control | 0 | 0 | 0 | 0 | 0 | 0 | Uninfected |
| 199 | M | Control | 0 | 0 | 0 | 0 | 0 | 0 | Uninfected |
| 206 | F | Control | 0 | 0 | 0 | 0 | 0 | 0 | Uninfected |
| 209 | F | Control | 0 | 0 | 0 | 0 | 0 | 0 | Uninfected |
| 211 | F | Control | 0 | 0 | 0 | 0 | 0 | 0 | Uninfected |
| 216 | M | Control | 0 | 0 | 0 | 0 | 0 | 0 | Uninfected |
| 224 | F | Control | 0 | 0 | 0 | 0 | 0 | 0 | Uninfected |
| 228 | F | Control | 0 | 0 | 0 | 0 | 0 | 0 | Uninfected |
| 230 | M | Control | 0 | 0 | 0 | 0 | 0 | 0 | Uninfected |
| 231 | M | Control | 0 | 0 | 0 | 0 | 0 | 0 | Uninfected |
| 192 | F | Infected | 3 | 73 | 0 | 0 | 140 | 1898 | Infected |
| 200 | F | Infected | 3 | 3040 | 0 | 0 | 3677 | 1780 | Infected |
| 202 | M | Infected | 3 | 46 | 0 | 0 | 15149 | 357 | Infected |
| 204 | M | Infected | 4 | 3 | 0 | 1 | 5539 | 160 | Infected |
| 205 | M | Infected | 0 | 0 | 0 | 0 | 0 | 0 | Infected |
| 214 | F | Infected | 2 | 0 | 0 | 0 | 89 | 138 | Infected |
| 217 | NA | Infected | 3 | 0 | 624 | 0 | 4498 | 278 | Infected |
| 218 | F | Infected | 4 | 912 | 5128 | 0 | 52521 | 570 | Infected |
| 220 | M | Infected | 4 | 23 | 122 | 0 | 485 | 370 | Infected |
| 222 | F | Infected | 4 | 27 | 947 | 0 | 19384 | 1941 | Infected |
| 223 | M | Infected | 0 | 0 | 0 | 0 | 0 | 0 | Uninfected[k] |
| 225 | M | Infected | 2 | 0 | 0 | 0 | 41119 | 83549 | Infected |
| 226 | M | Infected | 5 | 342 | 33 | 3 | 24691 | 27739 | Infected |
| 229 | M | Infected | 2 | 0 | 0 | 0 | 9 | 12 | Infected |

[a] Vole ID is the unique identification number assigned to each bank vole in the study.
[b] Sex refers to whether the bank vole is female (F) or male (M).
[c] Treatment refers to whether the bank vole was randomly assigned to the *B. afzelii*-infected group or the uninfected control group.
[d] N positive is the number of organs that tested positive for *B. afzelii* for each bank vole.
[e] Bladder is the number of spirochetes per mg of tissue in the bladder as estimated by our qPCR assay.
[f] Ear is the number of spirochetes per mg of tissue in the ear as estimated by our qPCR assay.
[g] Heart is the number of spirochetes per mg of tissue in the heart as estimated by our qPCR assay.
[h] Skin belly is the number of spirochetes per mg of tissue in the belly skin as estimated by our qPCR assay.
[i] Skin neck is the number of spirochetes per mg of tissue in the neck skin as estimated by our qPCR assay.
[j] Infection status is whether a vole was considered to be infected with *B. afzelii* or not (from Table S1).
[k] Bank vole 223 did not become infected following exposure to *B. afzelii*-infected ticks.

#### References

1. **Jacquet, M., J. Durand, O. Rais, and M. J. Voordouw.** 2015. Cross-reactive acquired immunity influences transmission success of the Lyme disease pathogen, *Borrelia afzelii*. Infection Genetics and Evolution **36:**131-140.
2. **Raberg, L.** 2012. Infection intensity and infectivity of the tick-borne pathogen *Borrelia afzelii*. J Evol Biol **25:**1448-1453.
3. **Gern, L., M. Siegenthaler, C. M. Hu, S. Leuba-Garcia, P. F. Humair, and J. Moret.** 1994. *Borrelia burgdorferi* in rodents (*Apodemus flavicollis* and *A. sylvaticus*): Duration and enhancement of infectivity for *Ixodes ricinus* ticks. Eur J Epidemiol **10:**75-80.
4. **Belli, A., A. Sarr, O. Rais, R. O. M. Rego, and M. J. Voordouw.** 2017. Ticks infected via co-feeding transmission can transmit Lyme borreliosis to vertebrate hosts. Scientific Reports **7:**5006.
5. **Jacquet, M., J. Durand, O. Rais, and M. J. Voordouw.** 2016. Strain-specific antibodies reduce co-feeding transmission of the Lyme disease pathogen, *Borrelia afzelii*. Environ Microbiol **18:**833-845.
6. **Rego, R. O. M., A. Bestor, J. Stefka, and P. A. Rosa.** 2014. Population bottlenecks during the infectious cycle of the Lyme disease spirochete *Borrelia burgdorferi*. PLOS ONE **9:**e101009.
7. **Durand, J., C. Herrmann, D. Genné, A. Sarr, L. Gern, and M. J. Voordouw.** 2017. Multistrain infections with Lyme borreliosis pathogens in the tick vector. Appl Environ Microbiol **83**.
